# Unscheduled DNA replication in G1 causes genome instability through head-to-tail replication fork collisions

**DOI:** 10.1101/2021.09.06.459115

**Authors:** Karl-Uwe Reusswig, Julia Bittmann, Martina Peritore, Michael Wierer, Matthias Mann, Boris Pfander

**Affiliations:** DNA Replication and Genome Integrity, Max Planck Institute of Biochemistry, Martinsried, 82152, Germany; Proteomics and Signal Transduction, Max Planck Institute of Biochemistry, Martinsried, 82152, Germany; Proteomics Research Infrastructure, University of Copenhagen, Copenhagen, 2200, Denmark

## Abstract

DNA replicates once per cell cycle. Interfering with the regulation of DNA replication initiation generates genome instability through over-replication and has been linked to early stages of cancer development. Here, we engineered genetic systems in budding yeast to induce unscheduled replication in the G1-phase of the cell cycle. Unscheduled G1 replication initiated at canonical S-phase origins across the genome. We quantified differences in replisomes in G1- and S-phase and identified firing factors, polymerase α, and histone supply as factors that limit replication outside S-phase. G1 replication per se did not trigger cellular checkpoints. Subsequent replication during S-phase, however, resulted in over-replication and led to chromosome breaks via head-to-tail replication fork collisions that are marked by chromosome-wide, strand-biased occurrence of RPA-bound single-stranded DNA. Low-level, sporadic induction of G1 replication induced an identical response, indicating findings from synthetic systems are applicable to naturally occurring scenarios of unscheduled replication initiation by G1/S deregulation.

## Introduction

To ensure that DNA is replicated precisely once per cell cycle, eukaryotic DNA replication initiation involves two steps, with each being restricted to different cell cycle phases (Bell and Labib, 2016). In the first step (origin licensing) replicative helicase precursors are loaded at origins of replication (Bleichert, 2019); in the second step (origin firing), cyclin-dependent kinase (CDK) and Dbf4-dependent kinase (DDK) promote helicase activation by facilitating the association of helicase accessory factors (Moiseeva and Bakkenist, 2018; Siddiqui et al., 2013; Tanaka and Araki, 2013). Multiple regulatory mechanisms ensure temporal separation of the two steps. Experimentally inducing unscheduled helicase loading or unscheduled helicase activation results in over-replication and genome instability (Green et al., 2006; Reußwig and Pfander, 2019; Reußwig et al., 2016; Tanny et al., 2006; Zhou et al., 2020).

Genome instability caused by unscheduled helicase loading has been linked to early stages of cancer development (Champeris Tsaniras et al., 2014; Mughal et al., 2019; Muñoz et al., 2017; Vassilev and DePamphilis, 2017). Unscheduled helicase activation is expected to have similar outcomes but currently there is no experimental system to test this hypothesis in human cells. It is therefore unknown whether unscheduled replication occurs upon de-regulation of the G1/S transition by oncogenes. Common oncogenic drivers, such as overexpression of cyclin E or MYC, cause oncogene-induced replication stress (Bartkova et al., 2006; Costantino et al., 2014; Macheret and Halazonetis, 2018) but current methodology is unable to determine if over-replication is occurring after oncogene induction. A marker for unscheduled replication would therefore facilitate the detection and investigation of early stages of cancer development.

Over-replication has mostly been studied in budding yeast via unscheduled helicase loading in the S/M phase of the cell cycle, induced by bypassing the cell cycle control of key licensing proteins (Nguyen and J. J. Li, 2001). CDK restricts helicase loading by a variety of mechanisms and unscheduled helicase loading is robustly blocked by several mechanisms that regulate licensing factors (Drury and Diffley, 2009; Green et al., 2006; Tanny et al., 2006). Abolishing the licensing control mechanisms induces repeated helicase loading and over-replication, resulting in replication fork collisions and double-strand breaks (DSBs) (Green et al., 2010; Green and J. J. Li, 2005), which leads to hallmarks of genome instability including gene amplifications, gross chromosomal rearrangements (GCRs) and aneuploidy (Archambault et al., 2005; Bui and J. J. Li, 2019; Finn and J. J. Li, 2013; Green et al., 2010; 2006; Green and J. J. Li, 2005; Hanlon and J. J. Li, 2015; Nguyen and J. J. Li, 2001; Tanny et al., 2006).

In metazoans, additional CDK-independent mechanisms regulate helicase loading and loss of replication control has also been investigated in these organisms (Arias and J. C. Walter, 2005; McGarry and Kirschner, 1998; Tada et al., 2001; Wohlschlegel et al., 2000). For example, unscheduled helicase loading results in extensive DNA damage and loss of cellular viability in cultured human cells and *Xenopus laevis* egg extracts (J. R. Hall et al., 2008; Jang et al., 2018; Klotz-Noack et al., 2012; Machida and Dutta, 2007; Maiorano et al., 2005; Melixetian et al., 2004; Neelsen et al., 2013; D. Walter et al., 2016; Zhu et al., 2004). In *Drosophila melanogaster*, follicle cells undergo developmentally programmed over-replication at specific genomic loci and DSBs occur at sites of potential head-to-tail replication-fork collisions (Alexander et al., 2015; 2016; J. C. Kim et al., 2011; Nordman et al., 2014). Thus, both natural and synthetic over-replication systems appear to generate DSBs and one can speculate that some form of fork stalling or collision is involved in generating this damage.

Unscheduled helicase activation could lead to replication in G1 or at the G1/S transition. However, our knowledge of its consequences is sparse, as currently experimental induction of G1 replication is only possible in budding yeast, where the minimal set of CDK targets has been identified to override cell cycle control of helicase activation (Tanaka and Araki, 2011; Tanaka et al., 2007; Zegerman and Diffley, 2007). Previous work indicated that unscheduled helicase activation results in genome instability, aneuploidy, and cell death (Tanaka and Araki, 2011), but little is known about how cells respond to over-replicated DNA in G1, what consequences this has for the following S-phase, and what kind of DNA structures are generated. Furthermore, it is unknown if there are other mechanisms constraining unscheduled replication in G1 besides the requirement for the activity of CDK and DDK.

Here, we engineered different genetic systems to induce unscheduled helicase activation and thereby replication in G1 in budding yeast and investigated its characteristics, constraints, and consequences. We found that unscheduled replication in G1 initiates at canonical origins on all chromosomes but progressed slower than canonical S-phase replication. Quantitative proteomics revealed a reduced number of replisomes, but also differences in replisome composition compared to S-phase replication. Testing for factors that constrain G1 replication we found histone availability to be limiting, suggesting that histone supply is a crucial bottleneck for replisome progression. Importantly, when we investigated the consequences of unscheduled G1 replication, we found that subsequent S-phase replication strongly aggravated genome instability. Specifically, we observed chromosome breaks and DNA damage checkpoint activation after release into S-phase. These phenotypes were completely suppressed when further replication initiation in S-phase was blocked indicating that successive rounds of replication caused the observed DNA damage. Data from strand-specific ChIP-sequencing of RPA-bound single-stranded DNA revealed a characteristic pattern of strand-biased RPA accumulation along whole chromosomes arising from single-ended DSBs that were generated by head-to-tail replication fork collisions and resected. Using a complementary strategy, we induced low levels of sporadic G1 replication and observed a similar cellular response indicating that our engineered systems reveal insights of physiological significance and that asymmetric accumulation of RPA-bound single-stranded DNA is a highly sensitive marker of acute over-replication.

## Results

### Unscheduled replication in G1 initiates at canonical replication origins

To engineer a system able to initiate unscheduled replication in G1, we adapted previously published strategies that allow the bypass of CDK and DDK control of replication (Tak et al., 2006; Tanaka et al., 2007; Zegerman and Diffley, 2007). In order to minimally interfere with cellular physiology, we implemented conditional expression of replication initiation proteins from galactose-inducible promoters (Fig. 1A). Expression of high levels of Dpb11 together with a CDK phosphorylation-mimicking allele of *SLD2* (*sld2-T84D*) generates a bypass to CDK-regulation of replication initiation and additional expression of the cell cycle-regulated DDK subunit Dbf4 allowed DDK activation in G1 (Cheng et al., 1999; Ferreira et al., 2000; Oshiro et al., 1999).

**Figure 1:**
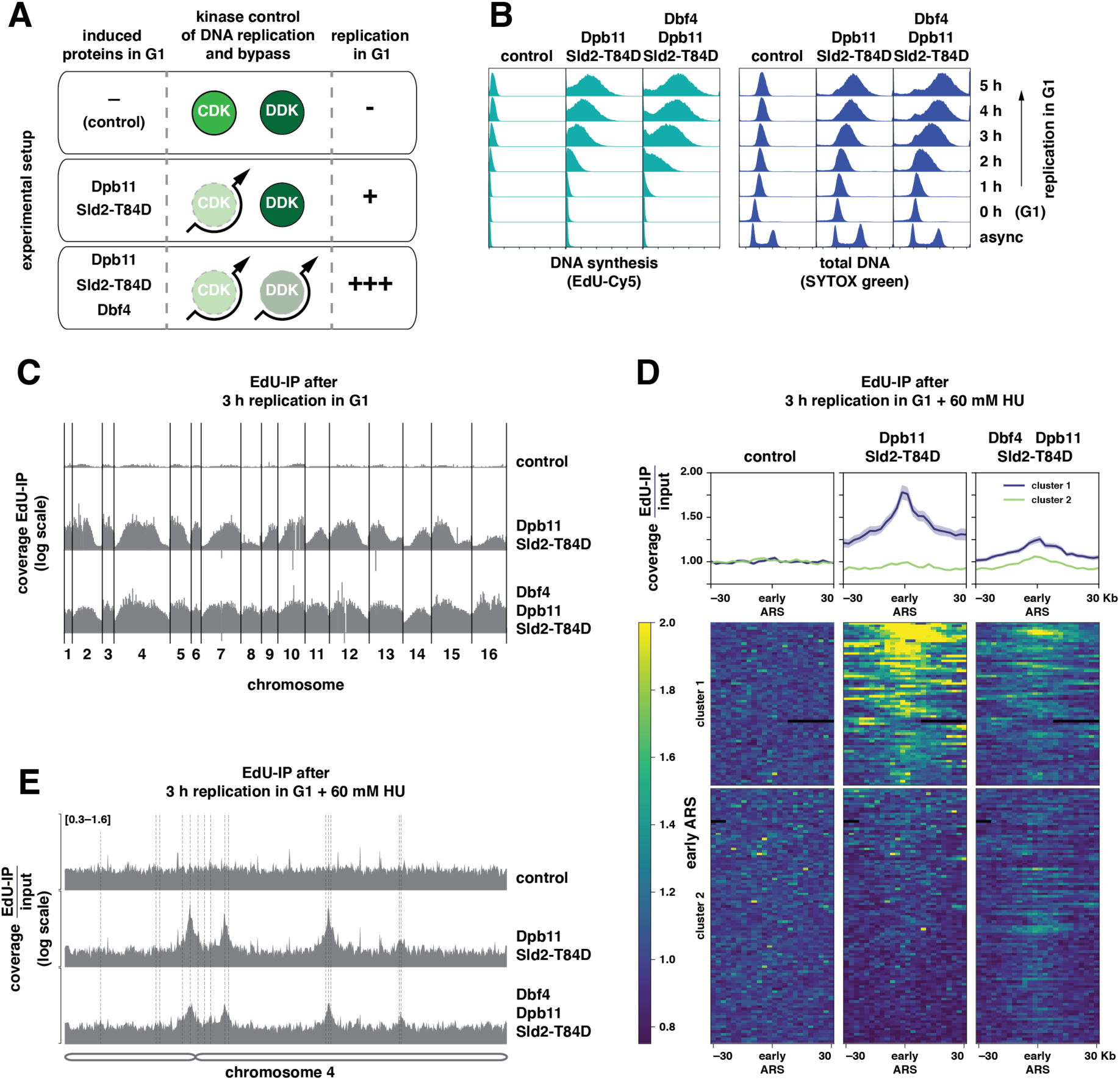
Unscheduled replication in G1 initiates at canonical replication origins genome-wide. (A) Summary of engineered genetic changes that allow to bypass cell cycle control of DNA replication and induce unscheduled replication in G1. Indicated proteins and variants are expressed from *pGAL1-10* promoter inducible by galactose. Experimental setup for G1 replication involves G1 cell cycle arrest using 10 µg/ml *α*-factor in 2% raffinose medium, followed by induction of G1 replication by 2% galactose. (B) Bypassing CDK control (Dpb11, Sld2-T84D) and CDK/DDK control (Dbf4, Dpb11, Sld2-T84D) generates different levels of unscheduled replication in G1. Cy5-labeled EdU incorporated after induction of replication in G1-arrested cells and SYTOX green-stained total DNA content were measured by flow cytometry at indicated timepoints. (C) Unscheduled replication in G1 after bypass of CDK or CDK/DDK control occurs genome-wide. EdU-labeled DNA as a proxy for DNA synthesis was isolated after 3 h of G1 replication and mapped to all sixteen *S. cerevisiae* chromosomes. Data from n=2 replicates. (D) and (E) Unscheduled G1 replication initiates at canonical replication origins. (D) Input-normalized coverage of 60 kb windows around early-replicating replication origins (autonomous replicating sequences (ARS)) with EdU-labeled DNA after 3 h replication in G1 in the presence of 60 mM hydroxyurea (HU) separated into two clusters based on mean signal intensity. (top) Profile plots of mean coverage (dark) ± SE (light). (bottom) Heatmaps with 2.5 Kb bin size. Data from n=2 replicates. (E) Representative EdU-IP coverage traces from the same experiment spanning the entire chromosome 4. Dotted lines indicate early-replicating ARSs. See also Figure S1.

We followed DNA synthesis in G1-arrested cells by flow cytometry measuring either incorporation of the nucleoside analog EdU during DNA synthesis or the increase in total cellular DNA (Fig. 1B). We observed a linear increase in DNA content over time, resulting in a 47% increase in average DNA content 5 h after induction with CDK bypass and in a 78% increase with CDK/DDK bypass compared to control cells that showed low levels of DNA synthesis arising from cell cycle-independent mitochondrial DNA replication (Fig. S1A). In all conditions, changes in total DNA correlated with changes in EdU-labeled DNA, indicating newly synthesized DNA was quantitatively labeled (Fig. S1B). Thus, both systems can be used to trigger unscheduled replication in G1 and tune the level of G1 replication.

To assess if unscheduled replication in G1 occurs genome-wide, we induced both systems for 3 h in G1-arrested cells and purified EdU-labeled DNA replication products via biotin handles for next-generation sequencing. Replication products were observed from all chromosomes, albeit to different extents (Fig. 1C): CDK bypass enriched for products from chromosomal regions that also replicate early in a regular S-phase (Fig. 1C middle, Fig. S1C), whereas bypass of both CDK and DDK regulation resulted in a more even coverage of all chromosomes, but late replicating regions and particularly those close to telomeres were still underrepresented in these samples (Fig. 1C bottom, Fig. S1C). Thus, unscheduled replication in G1 occurs in both systems to different extents on all chromosomes and appears to follow the same relative timing as replication in S-phase.

To determine if unscheduled G1 replication initiates from canonical replication origins, we limited DNA synthesis by addition of 60 mM hydroxyurea (HU) and purified EdU-labeled DNA replication products for next-generation sequencing (Fig. 1D, Fig. 1E). We detected replication initiation at origins which fire early in S-phase (Raghuraman et al., 2001), as indicated by symmetrical peaks around these loci. In the CDK bypass, DNA replication initiated from a subset of these early-firing origins (labeled ‘cluster 1’) while in CDK/DDK bypass conditions most early origins were used (Fig. 1D, Fig. 1E), consistent with replication occurring more evenly across chromosomes (Fig. 1C). Thus, unscheduled replication in G1 initiates from the same canonical replication origins as in S-phase. Furthermore, the differences between the CDK and CDK/DDK bypass indicate that CDK and DDK activation collectively leads to replication initiation from early-firing origins, while with limited DDK activation only a subset of these origins becomes active.

### G1 replisomes differ from S replisomes

We assessed whether replisome composition differs in G1-phase versus S-phase. To purify replisomes, we immunoprecipitated the GINS complex (Gambus et al., 2006), an integral part of the replicative CMG helicase (Cdc45–Mcm2-7–GINS), via a GFP-tag on its Psf2 subunit and measured replisome composition by label-free quantitative mass spectrometry. As a benchmark, we purified replisomes from untreated S-phase cells or HU-treated S-phase cells and compared them to untagged control strains. All replisomes had a protein composition consistent with previous studies (Fig. 2A, Fig. 2B) (Gambus et al., 2006). The abundance of individual replisome sub-complexes in the final purification varied substantially (Fig. 2D), allowing us to identify sub-complexes that either interact transiently during S-phase or dissociate from replisomes at different rates during purification (Fig. S2). When we compared replisomes from S-phase and HU-treated cells, we observed a twofold reduced abundance of CMG helicases in the HU sample (Fig. 2D), but additional association of the DNA repair protein Rad5 (Fig. 2A, Fig. 2B) which is known to act in response to replication fork stalling in HU-treated cells (Blastyák et al., 2007; Minca and Kowalski, 2010).

**Figure 2:**
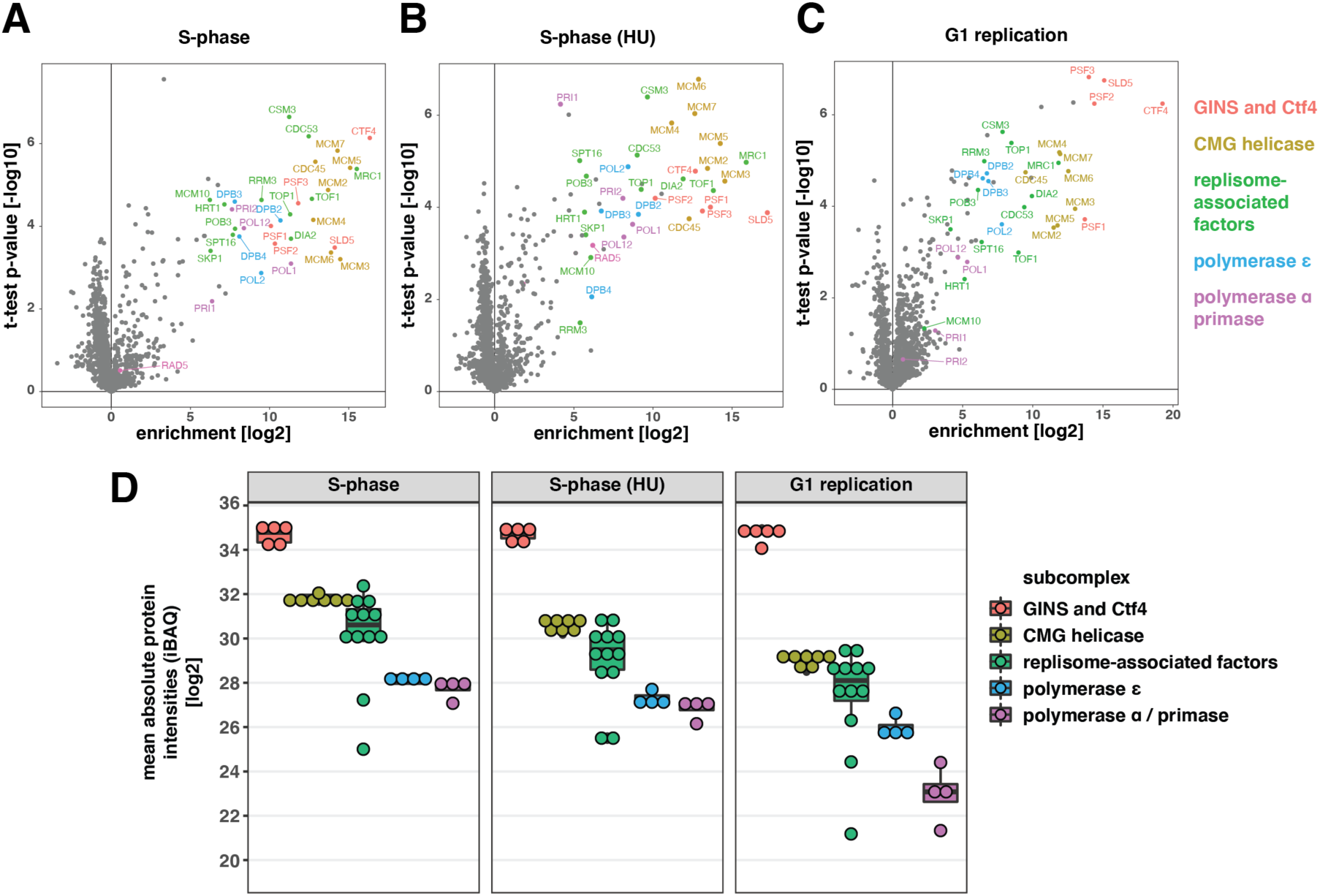
G1 replisomes differ quantitatively in subunit composition from S-phase replisomes. (A) to (C) Replisomes in S- and G1-phase contain the same set of proteins. Volcano-plot representation of mass-spectrometric analysis of replisomes by label-free quantification. Control cells were synchronously (A) released into S-phase, (B) arrested in S-phase using hydroxyurea (HU), or (C) G1 replication was induced for 3 h. Replisomes were purified via GFP-tagged GINS-subunit Psf2 and compared to untagged control samples. Colors indicate different replisome subcomplexes. Data from n=3 replicates per condition. (D) Reduced association of polymerase α/primase with replisomes in G1. Label-free quantification and comparison of the datasets shown in (A), (B), and (C) using intensity-based absolute quantification (iBAQ). See also Figure S2.

S-phase and G1 replication replisomes had qualitatively similar protein compositions (Fig. 2A, Fig. 2C), however the G1 sample had an eightfold reduction in assembled CMG, indicating the presence of fewer replisomes and therefore less efficient replication initiation. The leading strand DNA polymerase ε, fork protection complex Mrc1-Tof1-Csm3, topoisomerase Top1, helicase Rrm3, histone chaperone FACT, and ubiquitin ligase SCF-Dia2 all bound in similar relative ratios to both G1 and S replisomes (Fig. 2C, Fig. 2D). In contrast, the association of DNA polymerase α/primase and helicase activator Mcm10 with replisomes was reduced during G1 replication (Fig. 2C, Fig. 2D), suggesting the existence of another layer of cell cycle regulation acting at the step of replication priming and helicase activation. Thus, G1 replisomes have the same protein composition but form less efficiently and show reduced association of polymerase α and Mcm10.

### Factors limiting unscheduled G1 replication

To determine what constrains the formation of replisomes in G1 compared to S-phase we used our flow cytometry-based experimental setup. We considered the following factors could potentially restrict unscheduled replication in G1: depletion of licensed origins, ineffective bypass of CDK and/or DDK phosphorylation, low-abundance of firing factors in G1 (Mantiero et al., 2011; Tanaka et al., 2011), and insufficient supply of dNTPs as well as histones (Forey et al., 2020; Guarino et al., 2014; Marzluff and Duronio, 2002; Mendiratta et al., 2019).

To ask whether licensed origins may become depleted during G1 replication, we increased origin licensing activity by overexpressing the helicase loading factor Cdc6 which is tightly regulated through degradation at various cell cycle stages (Drury et al., 2000). Cdc6 overexpression did not affect the total amount of replication in G1 nor its initiation kinetics (Fig. 3A), indicating that the number of licensed origins in G1 cells does not limit replication in G1 in the CDK/DDK bypass setup.

**Figure 3:**
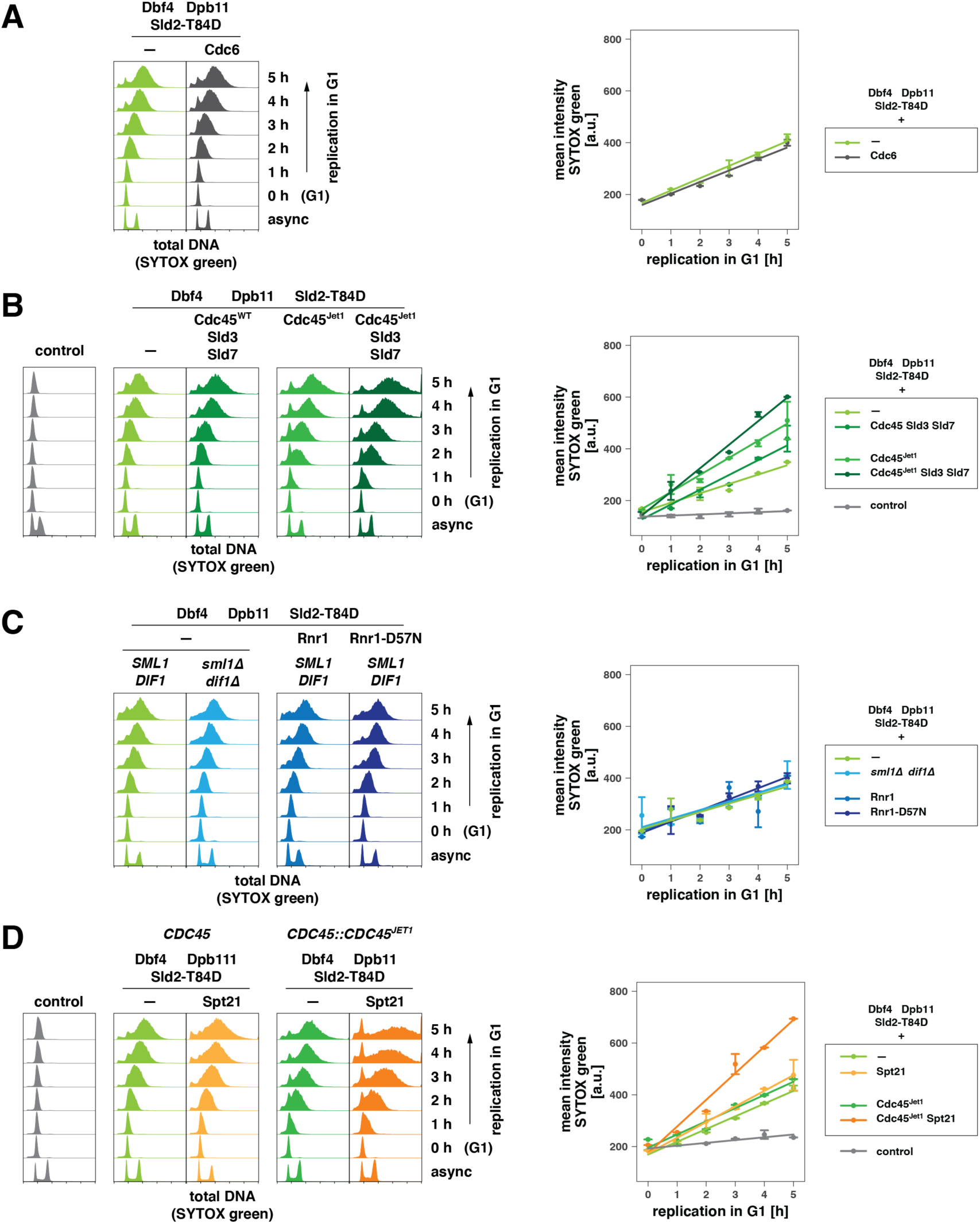
Factors limiting unscheduled G1 replication include availability of firing factors and histones. (A) Licensed origins do not limit levels of unscheduled G1 replication. Strains expressed licensing factor Cdc6 in addition to the CDK/DDK bypass system. (left) SYTOX green-stained total DNA after induction of G1 replication by galactose-induced expression of indicated proteins measured by flow cytometry at indicated timepoints. (right) Quantification of total DNA data in right panel by approximation of a bimodal distribution and calculating means of individual normal distributions. The average mean from 5 fits ± SD per timepoint is shown together with a linear regression. (B) CDK/DDK bypass for unscheduled replication in G1 is limited by the availability of initiation factors and efficient bypass of CDK-control. Experiment as in (A). (C) Activation of ribonucleotide reductase (RNR) does not lead to an increase of G1 replication. Experiment as in (A) using strains expressing the indicated *RNR1* alleles or lacking negative RNR regulators Sml1 and Dif1. (D) Increasing histone availability via expression of transcription factor Spt21 increases unscheduled replication in G1. Experiment as in (A). See also Figure S3.

Origin firing is known to be limited by the availability of replication initiation proteins (Mantiero et al., 2011; Tanaka et al., 2011). Therefore, in addition to Dbf4, Dpb11, and Sld2-T84D expression in G1-arrested cells, we also expressed Sld3, Sld7, and Cdc45 from a galactose-inducible promoter and observed a moderate increase in DNA synthesis compared to the basic CDK/DDK bypass strain (Fig. 3B). This suggests unscheduled G1 replication is constrained by the low abundance of firing factors.

The *CDC45^JET1^* allele is suggested to enhance binding of the Cdc45 protein to Sld3 and thereby bypass the requirement for CDK-phosphorylation of Sld3 (Tanaka et al., 2007; Tanaka and Araki, 2011). Cdc45^Jet1^ expression led to increased G1 replication, detectable after 1 h of induction when combined with the basic CDK/DDK bypass system (Fig. 3B). In contrast, deleting the DDK-antagonizing PP1-phosphatase targeting subunit *RIF1* (Davé et al., 2014; Hiraga et al., 2014; Mattarocci et al., 2014) or expressing an overactive, degradation-resistant allele of *DBF4* (*dbf4^RxxL-4A^*) (Cheng et al., 1999; Ferreira et al., 2000; Oshiro et al., 1999) did not alter the extent or the kinetics of unscheduled replication in G1 suggesting that DDK activity is not limiting in our setup (Fig. S3A). Taken together, further facilitating the bypass of CDK control of origin firing increases G1 replication, likely due to enhanced replication initiation.

Given levels of dNTPs and histone proteins rise at the G1-S transition to ensure effective genome replication (Guarino et al., 2014; Marzluff and Duronio, 2002), we tested whether availability of dNTPs and histones limits G1 replication. To increase dNTP concentrations in G1, we either deleted the ribonucleotide reductase inhibitors *SML1* and *DIF1* and/or over-expressed the catalytic subunit *RNR1* of ribonucleotide reductase as a wild-type or a D57N-allele, which is insensitive to feedback inhibition (Chabes et al., 1999; Chabes and Stillman, 2007; Lee et al., 2008; Zhao et al., 1998). Enhanced DNA synthesis was not observed in any of these conditions, suggesting that either concentrations of dNTPs are not a bottleneck to unscheduled replication in G1 or that additional G1-specific mechanisms exist, which suppress the rise of dNTP levels and affected our ability to experimentally induce dNTP synthesis in G1 (Fig. 3C).

Cell cycle regulation of the histone synthesis-promoting transcription factor Spt21 restricts expression of core histones to S-phase (Kurat et al., 2014). Ectopic expression of *SPT21* during G1 resulted in a marked increase in G1 replication induced by the CDK/DDK bypass system (Fig. 3D), indicating that lack of histone synthesis constitutes a bottleneck for unscheduled replication in G1. Moreover, expression of *SPT21* increased DNA synthesis synergistically with the *CDC45^JET1^* allele (Fig. 3D). Thus, efficient DNA replication can be reconstituted in G1 cells with major bottlenecks being an effective bypass of CDK control of origin firing and the low availability of histones in G1. These two factors could have complementary effects: While Cdc45^Jet1^ enhanced the efficiency of replication initiation as judged by the early increase of DNA content already 1 h after induction, Spt21 may improve replication elongation by promoting more efficient synthesis.

### G1-replication-induced DNA damage requires S-phase replication for checkpoint detection

To understand the consequences of unscheduled replication in G1 and how and when loss of replication control is detected, we induced unscheduled G1 replication and then released cells into the cell cycle. After G1 replication, cells entered and progressed through S-phase similar to control cells but subsequently entered cell cycle arrest suggesting DNA damage has occurred (Fig. S4A). Measurement of phosphorylated H2A (γH2A), a DNA damage marker, revealed that G1 replication induced low levels of γH2A in G1 (Fig. S4B), however passage through S-phase resulted in substantial accumulation of γH2A in late S/early M (Fig. 4A, Fig. 4B, 40 min after release). The γH2A increase was accompanied by the activation of the DNA damage checkpoint, as evidenced by hyper-phosphorylation of checkpoint kinase Rad53 (Fig. 4B). Checkpoint activation was dependent on the DNA damage-dependent checkpoint mediator Rad9, but not the replication checkpoint mediator Mrc1 (Fig. S4C, Fig. S4D). The checkpoint was not activated during G1 replication, even though replicating G1 cells were checkpoint-proficient (Fig. S4E, Fig. S4F). Thus, G1 replication per se does not trigger checkpoint activation, suggesting no widespread stalling of G1 replication forks.

**Figure 4:**
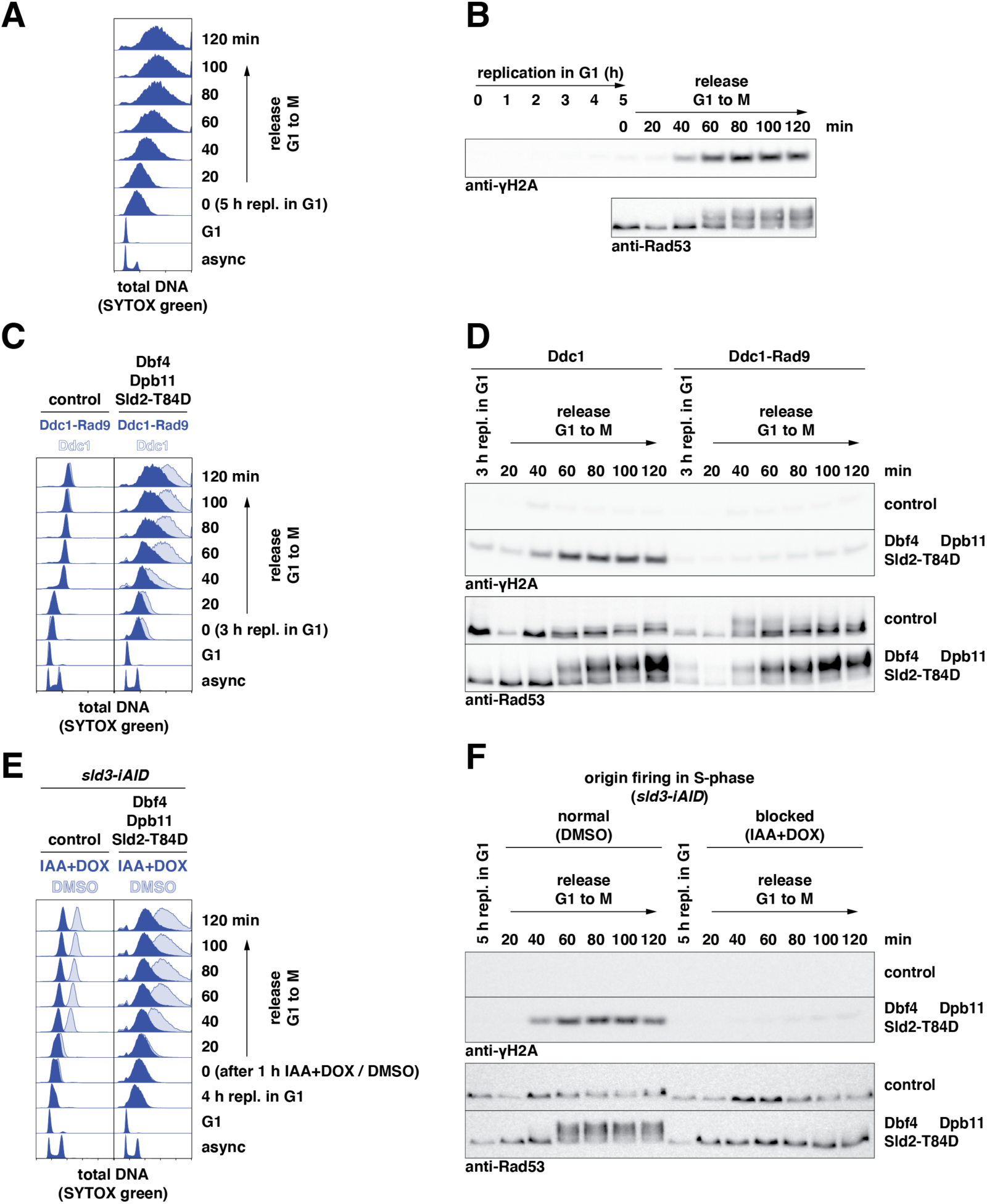
Unscheduled G1 replication induces DNA damage upon S-phase replication. (A) and (B) High levels of DNA damage and checkpoint activation occur late in the subsequent S-phase. To test the consequences of unscheduled G1 replication, G1 replication was induced for 5 h (CDK/DDK bypass), before cells were released from G1 arrest into the cell cycle (M phase arrest, nocodazole) and followed for indicated times. (A) SYTOX green-stained total DNA content as measured by flow cytometry at the indicated timepoints. (B) Western blots detecting levels of γH2A and Rad53, for which phosphorylated forms are visible by gel shift, at the indicated timepoints. (C) and (D) A hyper-sensitized DNA damage checkpoint restricts G1 replication and prevents DNA damage induction after release. Experiment as in (A)/(B) with strains expressing a Ddc1-Rad9 fusion protein as a second copy of Ddc1. (C) Flow cytometry data as in (A); (D) western blots as in (B). (E) and (F) Replication in S-phase is required for DNA damage induction and checkpoint activation. Experiment as in (A)/(B) with strains harboring a *sld3*-iAID degron allele to conditionally induce Sld3 degradation and suppress further origin firing. Depletion of Sld3 was triggered by addition of 3 mM auxin (IAA) and 20 µg/ml doxycycline (DOX) in the last hour of G1 replication before release. (E) Flow cytometry data as in (A); (F) western blots as in (B). See also Figure S4.

We hypothesized that the DNA damage checkpoint was not sufficiently sensitive to detect the low levels of damage signal arising from the small numbers of active replisomes operating during G1 replication as determined by mass spectrometry (Fig. 2D). To test this idea, we introduced the *DDC1-RAD9-fusion* allele, which increases the sensitivity of checkpoint signaling (Bantele et al., 2019; Pfander and Diffley, 2011), and observed that Rad53 was activated in response to G1 replication (Fig. 4C, Fig. 4D). These cells showed decreased amounts of DNA synthesis, both during G1 replication and the subsequent release (Fig. 4C), and the γH2A signal was largely suppressed (Fig. 4D). Thus, these cells activate a checkpoint-dependent block to origin firing (Lopez-Mosqueda et al., 2010; Santocanale and Diffley, 1998; Shirahige et al., 1998; Zegerman and Diffley, 2010), thereby limiting DNA synthesis and preventing DNA damage. These data suggest that checkpoint controls lack sufficient sensitivity to detect unscheduled replication in G1, but that a more sensitive checkpoint could prevent excessive unscheduled replication and the occurrence of DNA damage.

To test the hypothesis that DNA replication in S-phase following G1 replication gave rise to DNA damage, we conditionally depleted the firing factor Sld3 from cells (Fig. 4E) using an optimized auxin-inducible degron system (Morawska and Ulrich, 2013; Tanaka, 2021; Tanaka et al., 2015). Induction of G1 replication, followed by Sld3 degradation, and release into S-phase allowed us to shut off replication initiation in S-phase, as observed in both control cells and cells that had undergone G1 replication (Fig. 4E). Notably, suppressing replication initiation in S-phase also suppressed the occurrence of DNA damage and the activation of the DNA damage checkpoint (Fig. 4F), indicating that conflicts between G1 and S replication lead to the occurrence of DNA damage.

To determine whether over-replication, which involves re-licensing of origins, causes the observed DNA damage during G1 replication, we depleted the licensing factor Cdc6 during G1 replication, using a similar degron approach (Fig. S4G, Fig. S4H). Cdc6-depleted cells undergoing G1 replication synthesized less DNA (Fig. S4G, Fig. S4H) and had substantially reduced levels of DNA damage, as indicated by γH2A (Fig. S4H). Thus, over-replication, which occurs to some degree during G1 replication and then more extensively in the following S-phase, promotes the DNA damage associated with unscheduled replication in G1.

### Successive G1 and S replication generate single-ended DSBs

To visualize when and where successive G1 and S replication induced DNA damage, we analyzed chromosomes from a time course experiment using pulsed-field agarose gel electrophoresis (PFGE). Full-length chromosomes enter the PFGE gel, while the presence of replication forks or repair structures traps affected chromosomes in the loading slot. Using Southern blot probes against a marker locus (*TRP1*) present on chromosomes 4 and 7 in the analyzed strains, we observed that replication structures were only present during S-phase in control cells (20 min, Fig. 5A). In contrast, chromosomes were largely retained in the loading slots if cells had previously undergone replication in G1 (Fig. 5A, Fig. S5A). The level of retention correlated with the amount of replication induced in G1, when comparing CDK bypass with CDK/DDK bypass conditions (Fig. 5A, Fig. S5A). We detected additional signals (smears) below the chromosome bands after 80 min of release, indicative of DNA double-strand breaks (DSBs), demonstrating that DSBs only occur at late timepoints after successive G1 and S replication.

**Figure 5:**
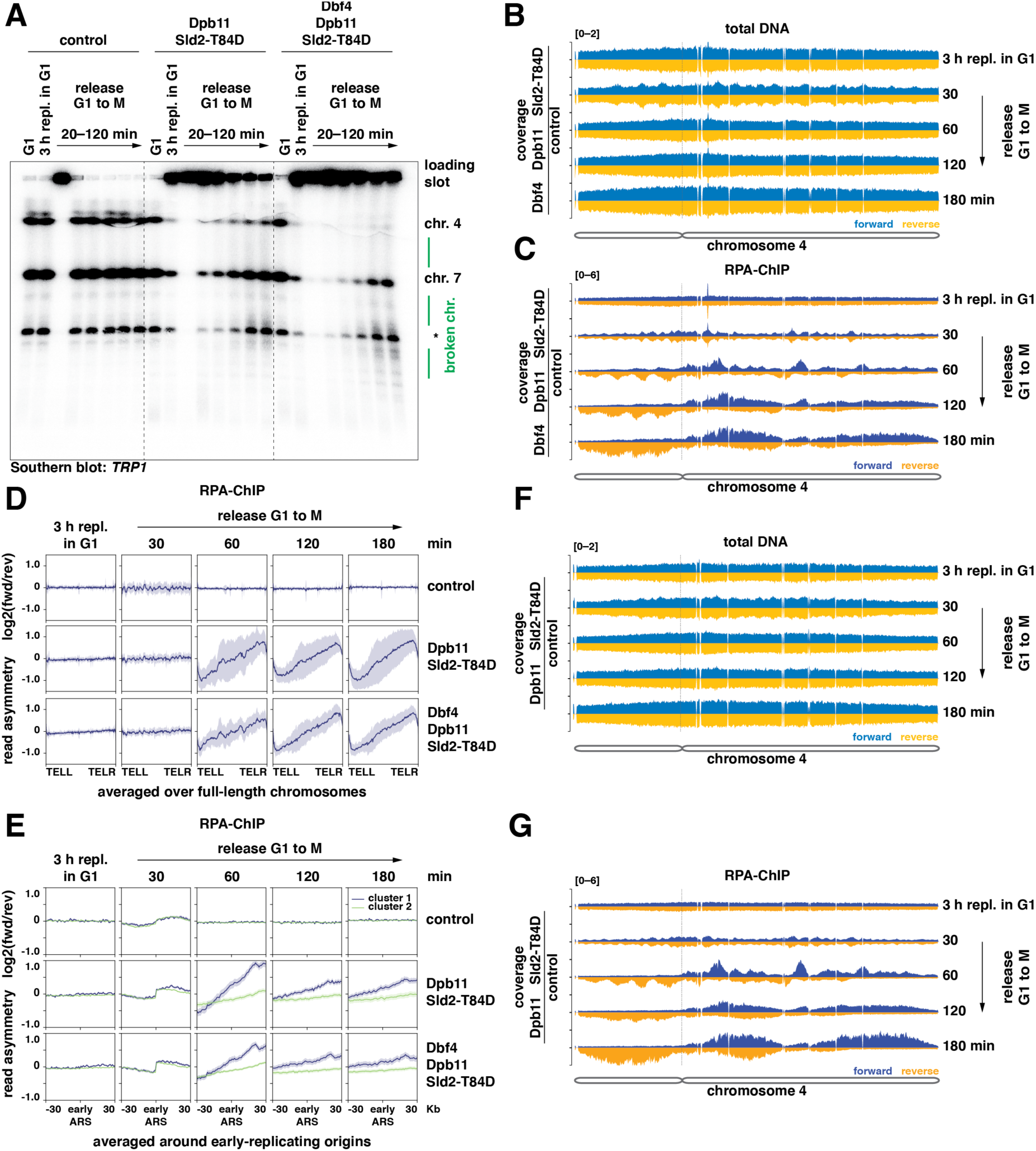
Successive G1 and S replication generate single-ended DSBs from head-to-tail fork collisions, resulting in an asymmetric pattern of RPA-bound ssDNA on chromosome arms. (A) Chromosome breaks occur after release from unscheduled G1 replication. Samples were taken at the indicated timepoints and chromosomes were separated by pulsed-field gel electrophoresis. A probe directed against the *TRP1* gene was used to visualize corresponding loci on chromosomes 4 and 7. Asterisk indicates an unspecific band. (B) and (C) RPA accumulates on chromosome arms with a strand bias. Representative traces of total DNA and RPA-ChIP from chromosome 4. G1 replication was induced by CDK/DDK bypass and cells were released to nocodazole-containing medium for the indicated times. Strand-specific forward reads are shown in light blue/dark blue; reverse reads are shown in yellow/orange. Dashed lines indicate the position of the centromere. Data from n=2 replicates. (D) and (E) RPA accumulates on chromosome arms in an asymmetric, characteristic pattern. The log2-ratio of RPA-ChIP-seq reads mapping to forward and reverse strands from experiment in (B)/(C) was averaged over full-length chromosomes (D) or a 60 Kb window around early replication origins (E) at indicated timepoints after release from unscheduled replication in G1 mediated by expression of the indicated set of proteins. Data are mean log2 ratio ± SD from n=2 replicates. (F) and (G) as in (B)/(C) but G1 replication was induced by CDK bypass. See also Figure S5.

DSBs arising from replication fork collisions will be either single-ended or double-ended. To distinguish between these two possibilities, and assuming that DSBs become resected, we used a ChIP-seq approach to study the strand-specific binding of RPA to single-stranded DNA (Peritore et al., 2021). We observed over-representation of regions around centromeres in total DNA after 3 h of G1 replication and subsequent release when comparing to a control strain, indicating preferential replication of these regions (Fig. S5B), consistent with our EdU-sequencing data (Fig. 1C). Regions of single-stranded DNA on chromosomes, as marked by increased RPA binding, appeared only after release from G1 arrest (Fig. S5C). RPA preferentially bound to the forward-strand DNA on the right arm of the chromosomes and to the reverse-strand DNA on the left arm of the chromosome (Fig. S5D). Both short and long chromosomes were similarly affected, as shown by RPA read asymmetry scores normalized for chromosome length (Fig. S5E). This asymmetric binding pattern was independent of *RAD52* (Fig. S5D), indicating that it does not involve recombination processes such as break-induced replication (BIR) (Davis and Symington, 2004; Ira and Haber, 2002; Kramara et al., 2018).

To analyze the appearance of the ssDNA-RPA signals, we carried out a time course experiment involving G1 replication induction for 3 h and then samples being taken at 0, 30, 60, 120, and 180 min following release into S-phase. Chromosome 4 (representative of all chromosomes) showed an over-representation of regions around centromeres in total DNA samples after G1 replication, demonstrating again that G1 replication is biased to centers of chromosomes (Fig. 5B). With cells in S-phase (30 min after release), we observed an over-representation of sequences close to origins of replication in total DNA (Fig. 5B) and an RPA pattern (Fig. 5C, Fig. 5D, Fig. 5E) consistent with single-stranded lagging strand template DNA (Fig. S5F, Fig. S5G). Averaging RPA-ChIP-seq data over all yeast chromosomes, we did not detect any chromosome-wide strand-biased RPA binding in control cells, after 3 h of G1 replication, nor in the first 30 min of the following S-phase (Fig. 5D).

However, 60 min after release and at later time points, a strand-biased RPA-ssDNA binding pattern developed over entire chromosomes (Fig. 5C, Fig. 5D). G1 replication by CDK bypass compared to G1 replication by CDK/DDK bypass led to RPA signals in the same chromosomal regions after the following S-phase, but with more pronounced strand bias throughout the genome (Fig. 5F, Fig. 5G, Fig. S5H). We reasoned that such chromosome-wide RPA-ssDNA strand bias would be generated if single-ended DSBs occur with a biased orientation. In our experiments, G1 replication preferentially initiated around centromeric regions of chromosomes (Fig. 1C) and traveled towards chromosome ends, recapitulating the inherent distribution of early- and late-replicating origins along chromosomes. Further replication initiation in S-phase then caused fork collisions that gave rise to single-ended DSBs accompanied by exposure of RPA-covered ssDNA on the forward or reverse strand depending on which direction the replication fork was moving. Focusing our analysis on origins which are active during G1 replication (Fig. 5E), our data supports a model (Fig. 7) where stochastic head-to-tail replication fork collisions between G1 replication forks and tailgating S-phase forks occur with directional bias towards chromosome ends and generate the observed RPA binding-patterns.

### Sporadic G1 replication generates fork collisions and genome instability

Given the increase in RPA-bound ssDNA did not scale with the amount of unscheduled G1 replication, we asked whether a single or few sporadic events of unscheduled replication initiation per cell could trigger similar cellular responses and genome instability. To test this idea, we devised an experimental setup to trigger sporadic, unscheduled replication in G1 by enhancing the physical interaction between firing factors Dpb11 and Sld2. We fused various split-Venus tags with Dpb11 and Sld2 proteins (Fig. S6A), which were expressed at similar levels but stabilized the interaction to different degrees (Fig. 6A, Fig. S6B). The combination of Dpb11-VN and VC-Sld2 yielded the highest Venus fluorescence intensity indicating that it stabilized the physical interaction most effectively (Fig. 6A, Fig. S6B). We assessed the extent of sporadic G1 replication by arresting cells in G1 in the presence of EdU and measured DNA synthesis at various times. We found that DNA replication initiated only in a sub-population of cells and relatively little DNA was replicated in these cells compared to the inducible G1 replication systems used before (Fig. S6C, compare to Fig. 1B). The combination of Dpb11-VN and VC-Sld2 yielded the highest level of replication in G1, consistent with the interaction data (Fig. 6A, Fig. S6B). Thus, sporadic initiation of DNA replication in G1 can be mediated by a synthetic Venus-bridged Dpb11-Sld2 complex.

**Figure 6:**
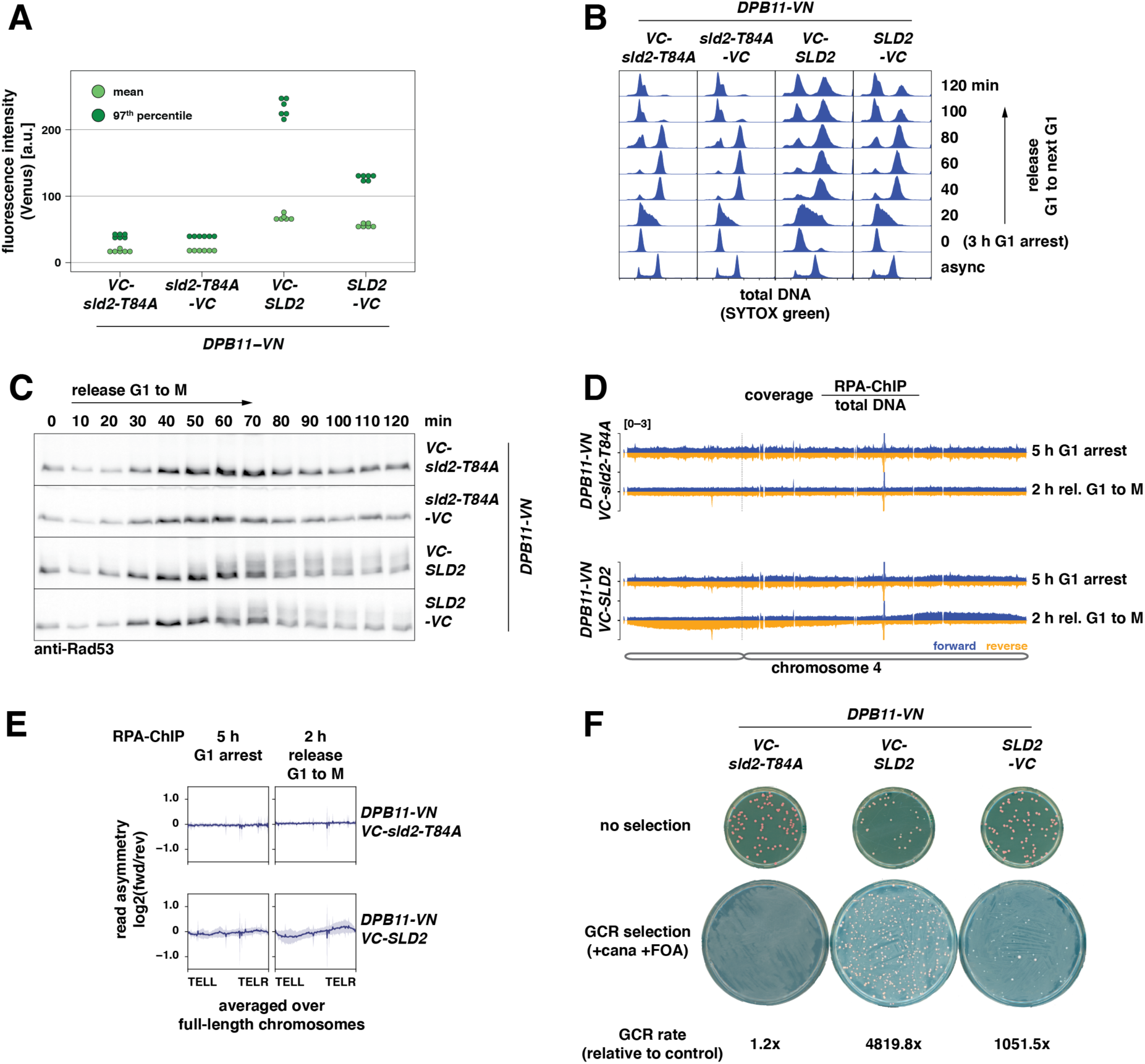
Low levels of sporadic G1 replication also generate head-to-tail fork collisions and genome instability. (A) Split-Venus tags (VN/VC) stabilize the physical interaction between Dpb11 and Sld2. Fluorescence intensity of strains expressing Dpb11-VN and Sld2 tagged at either N- or C-terminus with split-Venus fragment VC. Data represents mean (light green) and 97^th^ percentile (dark green) of split-Venus fluorescence intensity measured by flow cytometry in log-phase cells from n=6 replicates. Other combinations are shown in Fig. S6B. (B) Venus-stabilized interaction of Dpb11 and Sld2 results in cell cycle arrest. SYTOX green-stained total DNA from samples at indicated timepoints after release from G1 arrest to next G1-phase as measured by flow cytometry. (C) Venus-stabilized interaction of Dpb11 and Sld2 causes checkpoint activation after S-phase. Western blots detecting Rad53 and phosphorylated isoforms at indicated timepoints after G1 release from samples in (B). (D) and (E) Venus-stabilized interaction of Dpb11 and Sld2 results in asymmetric, strand-biased RPA binding on chromosome arms. (D) Coverage of the forward and reverse strand of chromosome 4 are shown for strains with Venus-stabilized Dpb11-Sld2 interaction *(VC-SLD2)* and interaction-deficient control (*VC-sld2-T84A*). Dashed line indicates the position of the centromere. (E) Log2-ratio of RPA-ChIP-seq reads mapping to forward and reverse strand was averaged over full-length chromosomes at indicated timepoints. Data are mean log2-ratio ± SD from n=2 replicates. (F) High levels of genome instability are caused by Venus-stabilized interaction of Dpb11 and Sld2. Representative control and selection plates from a standard assay detecting gross chromosomal rearrangements (GCRs). Absolute GCR rates are given in Fig. S6G. See also Figure S6.

To test if this sporadic system recapitulates the hallmarks of unscheduled G1 replication, we arrested cells in G1 and subsequently followed them through one round of the cell cycle until the next G1-phase (Fig. 6B, Fig. S6D). We detected phosphorylated Rad53 at 50-60 min after release (Fig. 6C), the level correlating with the amount of replication in G1, as shown by comparison of the most effective strain expressing VC-Sld2 to the less effective Sld2-VC strain (Fig. 6B, Fig. 6C). Consistent with this, G1 replication triggered by the sporadic system also resulted in cell cycle arrest (Fig. 6B, Fig. S6D) similar to strains after G1 replication by CDK/DDK bypass (Fig. S4A) with the number of arrested cells being proportional to the amount of G1 replication across different strains (Fig. S6C). Thus, sporadic replication in G1 occurs in a sub-population of cells that contains high levels of Venus-bridged Dpb11-Sld2 and results in checkpoint activation and cell cycle arrest after S-phase.

To assess if the sporadic system also leads to strand-biased detection of RPA-bound ssDNA on chromosome arms we conducted a strand-specific RPA-ChIP-seq experiment where we arrested cells for 5 h in G1 and then released them for 2 h to M phase. While we did not observe strand-biased RPA binding in control cells expressing an interaction deficient *VC-sld2-T84A* allele, we found that RPA bound preferentially to the forward strand on the right arm of chromosome 4 and to the reverse strand on its left arm (Fig. 6D, Fig. 6E). Such asymmetry was only detected on long yeast chromosomes that also contain many origins (Fig. S6E, Fig. S6F), suggesting that origin-rich chromosomes are more likely to engage in replication in G1 induced by the sporadic system and that G1 replication is a rare event in this system. This notion is consistent with the limited amount of DNA synthesis measured during G1 (Fig. S6C). Thus, strand-biased RPA binding on chromosome arms can be observed under conditions where only one or few origins initiate in an unscheduled manner during G1.

To determine if and how different levels of sporadic induction of unscheduled G1 replication cause genome instability, we selected strains expressing VC-tagged Sld2 (*VC-SLD2* and *SLD2-VC*) as they showed different levels of Dpb11-Sld2-complex formation and replication in G1 (Fig. 6A, Fig. S6C). Cultures of these strains were grown from single cells to saturation to determine gross chromosomal rearrangement (GCR) rates using an established assay (Putnam and Kolodner, 2010). GCRs are potent drivers of genome instability and frequently observed in cancer cells. We measured a highly increased GCR rate for the *SLD2-VC* (∼1000-fold compared to control) strain and an even higher GCR rate for the *VC-SLD2* (∼5000-fold) strain (Fig. 6F, Fig. S6G) suggesting that levels of genome instability correlated with the amount of sporadic G1 replication. Furthermore, cultures with increased levels of sporadic replication in G1 (*VC-SLD2*) showed decreased viability on non-selective medium, whereas cultures with lower levels of sporadic replication in G1 (*SLD2-VC*) had normal viability (Fig. 6F). Thus, our data suggest that unscheduled G1 replication induces genome instability and cell death, even when only single or few replication origins per cell are affected.

## Discussion

Over-replication has been linked to early stages of carcinogenesis (Petropoulos et al., 2019) but whether it is a cancer driver remains to determined. Previous studies in yeast and human cell lines have focused on mis-regulation of helicase loading factors and the induction of re-replication after S-phase (Nguyen and J. J. Li, 2001; Petropoulos et al., 2019). In contrast, many oncogenes act by de-regulating the G1-S transition, raising the potential of unscheduled DNA replication in G1 or early S-phase. Here, we induced unscheduled G1 replication in engineered budding yeast systems to reveal details of the molecular mechanism and cellular consequences of this toxic process.

We found that unscheduled helicase activation in G1 induced replication from canonical origins on all chromosomes with re-initiation at single origins being a rare event during G1 (Fig. 1C). Early-replicating origins are prone to both G1 replication and over-replication in our assays (Fig. 1D). The non-random distribution of over-replicated DNA was also observed in a recent study, implying that specific origins tend to participate in over-replication (Menzel et al., 2020). Such a preference has also been observed in cancer cells exposed to an experimental therapeutic strategy that induces overt over-replication (Fu et al., 2021). Over-replication induced by unscheduled helicase activation in G1 appears to differ in this regard from over-replication induced by unscheduled helicase loading in M phase, which re-initiates from a different set of replication origins that are flanked by specific re-initiation promoting sequence elements (Richardson and J. J. Li, 2014). This difference not only shows that regulated helicase loading is crucial for the establishment of the replication program, it also highlights that we cannot easily extrapolate findings from previous systems that induce re-replication after S-phase to unscheduled DNA replication in G1.

When compared to replication in S-phase, replication in G1 progressed approximately tenfold slower in bulk. As similar observations were made for systems of unscheduled replication in M phase (Nguyen and J. J. Li, 2001), we reasoned that additional factors may be constraining replication outside S-phase. We show that inefficient replication initiation and inefficient replication elongation both contribute to overall slow replication (Fig. 3). For replication elongation, we show that promoting histone synthesis in G1, which is normally a key feature of S-phase (Marzluff and Duronio, 2002), accelerated G1 replication by approximately twofold (Fig. 3D), suggesting that histone protein availability is a major bottleneck to replication in G1, even though repression of histone synthesis has only minor effects on S-phase length in budding yeast (U. J. Kim et al., 1988). Our mass spectrometry-based quantification revealed a reduced association of DNA polymerase α/primase with G1 replisomes (Fig. 2D). Polymerase α/primase has been proposed to be cell-cycle regulated (Foiani et al., 1997; Schub et al., 2001) and was found to be phosphorylated by CDK (Holt et al., 2009). Such phosphorylation could regulate its association with the replisome and, indeed, the efficiency of replication initiation in S-phase is decreased if protein levels of polymerase α fall below a threshold (Porcella et al., 2020). It is unclear whether cell cycle control of polymerase α/primase would influence primarily replication initiation or elongation or both, but nonetheless we demonstrate that studying unscheduled replication in G1 facilitates the identification of cell cycle control circuits.

Our study addresses the question of how cells respond to unscheduled replication and how G1 replication induces DNA damage. We show that G1 replication compromises genome stability and it does so specifically due to conflicts of G1 replication forks with subsequently initiated S replication forks (Fig. 4E, Fig. 4F). The initial G1 replication is not detected by cellular checkpoint controls, likely because it is carried out only by relatively few replisomes. Therefore, cells commence their “normal” DNA replication program during S-phase, during/after which high levels of DNA damage occur. We conclude a model (Fig. 7), whereby head-to-tail collisions of DNA replication forks are central to this DNA damage induction. G1 replication will leave behind replication forks (Fig. 7A) and initiation of replication in S-phase will generate new replication forks that have the propensity to tailgate onto G1 replication forks (Fig. 7B and Fig. 7C), no matter which strand is being replicated. Head-to-tail collisions of G1 and S replication forks will generate single-ended DSBs (Fig. 5A, Fig. 5D), which subsequently become subject to DNA end resection explaining the strand-biased appearance of single-stranded DNA (Fig. 5D). The mechanism of DSB induction through head-to-tail collisions of replication forks is thus similar to what has been proposed for over-replication induced by unscheduled helicase loading (Davidson et al., 2006; Green et al., 2010) and blocked replisome progression (Alexander et al., 2015). These collisions will initially be avoided (but not ultimately prevented) if the G1 replication fork encounters a fork in head-to-head orientation (Fig. 7D, Fig. 7E), leading to termination before the tailgating fork arrives. In this case, however, a new problem arises, because the tailgating S-phase fork will now remain unterminated, because it lacks a termination “partner” and therefore can lead to a head-to-tail collision with a neighboring S-phase fork. Due to the bi-directional nature of DNA replication, we think the only theoretical solution to this problem is re-duplication of the entire chromosome.

**Figure 7:**
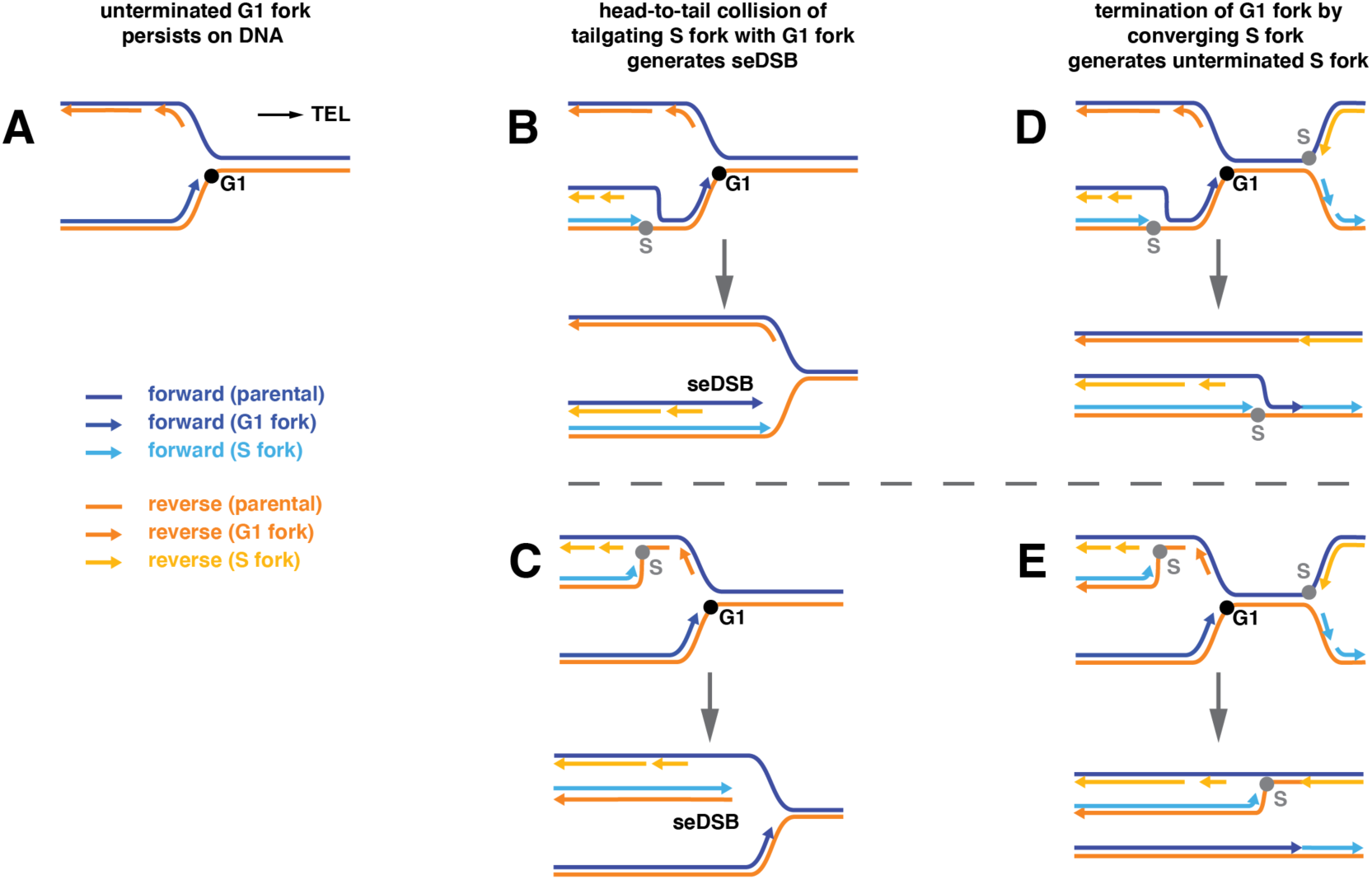
Single-ended double-strand breaks result from head-to-tail collisions of S forks with unterminated G1 forks. (A) Unterminated forks persist after unscheduled replication in G1 due to incomplete duplication. For simplification a single fork moving to the right telomere is shown. Since central regions are preferentially replicated during G1 replication, these forks are most frequently moving away outwards towards telomeres (TEL). (B) and (C) Head-to-tail collision of an S fork with an unterminated G1 fork. A single-ended double-strand break (seDSB) occurs independent of whether the S fork travels on the parental reverse strand (B) or the reverse strand synthesized by the G1 fork (C). (D) and (E) The G1 fork is terminated by a converging S fork, but this results in an unterminated S fork. Different scenarios separated as in (B)/(C).

Where collisions occur in the genome is determined by the location of the G1 replication fork relative to its two nearest origins, as well as their respective initiation timing. Because helicase activation is itself a stochastic process (Hennion et al., 2020; Müller et al., 2019; Wang et al., 2021), forks will also be resolved stochastically. We observed that G1 replication mimics early S-phase (Natsume et al., 2013) with replication initiating primarily from origins in central regions of chromosomes, including centromeres, but not towards chromosome ends (Fig. 1C). Therefore, G1 replication forks that will be involved in head-to-tail collisions will mainly be moving outwards towards telomeres. It is thus the timing and the relative efficiency of replication origins that shape where over-replication generates single-ended DSBs within the genome.

At this point we cannot exclude that replication run-off contributes to the occurrence of single-ended DSBs and single-stranded DNA. Such run-off will occur if the product of G1 replication, which is used as the template for S-phase replication, contains DNA nicks (single-strand breaks) or gaps. Indeed, large RPA-coated ssDNA gaps have been observed on the template strand during over-replication in human cells (Neelsen et al., 2013). In contrast, we do not observe evidence for the occurrence of large ssDNA gaps during G1 replication, but cannot exclude a contribution of DNA nicks (Vrtis et al., 2021).

Over-replication from multiple origins may be a rare event given the various endogenous replication control mechanisms. Sporadic unscheduled replication events affecting only single chromosomes may however be more likely, particularly under conditions of deregulated cell cycle control. Our model suggests that even a single event of unscheduled replication will be detrimental to the affected chromosome and can only be resolved if the entire chromosome is re-duplicated. To test this hypothesis, we generated systems to sporadically induce unscheduled replication in G1 (Fig. 6) and observed the same signature of asymmetric, strand-biased RPA accumulation (Fig. 6D), but only on long chromosomes (Fig. S6E), which we speculate undergo unscheduled replication more frequently due to the higher number of (early-firing) origins (Fig. S6F). We also observed a substantial increase in chromosomal rearrangements (Fig. 6F), highlighting the potential of rare over-replication events as potent drivers of genome instability.

Taken together, our analysis of unscheduled G1 replication revealed a characteristic pattern of chromosome-wide asymmetry of single-stranded DNA and associated RPA, which has the potential to serve as a marker for acute over-replication, be useful in clarifying the exact nature of replication stress, and help reveal the contribution of unscheduled replication to early carcinogenesis.

## Acknowledgments

We thank Uschi Schkölziger and Sandra Mitzkus for excellent technical assistance, Helle Ulrich, Etienne Schwob and Seiji Tanaka for yeast strains and plasmids, Jonathan Baxter, Armelle Lengronne, Philippe Pasero, Iestyn Whitehouse for protocols and/or experimental advice, Marja Driessen and Rin ho Kim (MPIB NGS core facility) for next-generation sequencing, Stefan Pettera (MPIB core facility) for peptide synthesis, Guy Riddihough and Life Science Editors for editing the manuscript, Dominik Boos, Stephan Hamperl, Christoph Kurat and all members of the Pfander lab for stimulating discussion and critical reading of the manuscript. This work was supported by the Max Planck Society (to BP, to MM) and grants by the Deutsche Forschungsgemeinschaft (DFG, German Research Foundation): PFA794-5/1 (to BP); Project-ID 213249687 – SFB 1064 (A23 to BP).

## Author contributions

K.-U.R. and B.P. conceived and designed the study. K.-U.R. and J.B. conducted experiments, M.P. prepared libraries for strand-specific RPA-ChIP-seq, M.W. measured and analyzed mass spectrometry data, K.-U.R. analyzed NGS data. K.-U.R. and B.P. wrote the manuscript. All authors analyzed data and commented on the manuscript. M.M. and B.P secured funding.

## Methods

### Yeast strains and culture

All yeast strains were constructed in the W303 background using standard methods (Janke et al., 2004). Genotypes of all used strains are given below (Table S1) and if not stated otherwise strains were constructed in the EdU-incorporating background of E3087. Integrative plasmids were linearized prior to transformation and single integration of plasmids was confirmed by PCR. Gene deletions and tags were introduced using a PCR-based protocol.

For cell cycle experiments, cells were grown to log-phase (OD600 of 0.5-0.6) at 30 °C in YP medium supplemented with adenine and either 2% raffinose (inducible G1 replication system) or 2% glucose (sporadic G1 replication system) and synchronized in G1 by adding α-factor (MPIB core facility or GenScript RP01002) to a final concentration of 0.5 µg/ml for *bar1Δ* cells or 10 µg/ml for *BAR1* cells. Additional doses of α-factor were added after each hour of arrest to achieve a stable arrest of *BAR1* cells. Hydroxyurea (Sigma H8627) was added to a final concentration of 200 mM to achieve an arrest in S-phase; nocodazole (Sigma M1404) was added to a final concentration of 5 µg/ml to achieve an arrest in M phase. Cell cycle arrest was confirmed by using a microscope and by taking samples for flow cytometry. To release cells from a cell cycle arrest, cells were washed once with and then re-suspended in pre-warmed YP-medium containing the appropriate sugar. To deplete cells of a protein carrying an auxin-inducible degron (AID) tag, indole-3-acetic acid (IAA, Sigma I3750) was added to 3 mM final concentration. A pre-treatment of cells with doxycycline was required to allow effective depletion via the iAID-system (Tanaka et al., 2015). Specifically, *sld3-iAID* cells were cultured in the presence of 0.1 µg/ml doxycycline (DOX, Sigma D9891) and the doxycycline concentration was increased to 20 µg/ml when IAA was added. EdU (Santa Cruz Biotechnology sc-284628) was used at a final concentration of 100 µM to label newly synthesized DNA.

**Table S1:** Budding yeast strains used in this study.

| strain | genotype | source / reference |
| --- | --- | --- |
| W303-1A | *MATa ade2-1 ura3-1 his3-11,15 trp1-1 leu2-3,112 can1-100* | (Rothstein, 1983) |
| E3087 | W303-1A *RAD5+*<br>*ura3::URA3/pGPD-TK(5x)*<br>*AUR1c::pADH-hENT1* | (Talarek et al., 2015) |
| YKR1445 | *leu2::pGAL-DPB11/sld2-T84D::LEU2* | this study |
| YKR1447 | *his3::pGAL-DBF4::HIS3*<br>*leu2::pGAL-DPB11/sld2-T84D::LEU2* | this study |
| YKR1500 | *AUR1c::pADH-hENT1*<br>*bar1Δ::natNT2* | this study |
| YKR1501 | *leu2::pGAL-DPB11/SLD2::LEU2*<br>*bar1Δ::natNT2* | this study |
| YKR1502 | *leu2::pGAL-DPB11/sld2-T84D::LEU2*<br>*bar1Δ::natNT2* | this study |
| YKR1503 | *his3::pGAL-DBF4::HIS3*<br>*leu2::pGAL-DPB11/sld2-T84D::LEU2*<br>*bar1Δ::natNT2* | this study |
| YKR1546 | *pep4Δ::kanMX4* | this study |
| YKR1558 | *pep4Δ::kanMX4*<br>*PSF2-yeGFP::hphNT1* | this study |
| YKR1553 | *pep4Δ::kanxM4*<br>*his3::pGAL-DBF4::HIS3*<br>*leu2::pGAL-DPB11/sld2-T84D::LEU2*<br>*bar1Δ::natNT2* | this study |
| YKR1557 | *pep4Δ::kanxM4*<br>*his3::pGAL-DBF4::HIS3*<br>*leu2::pGAL-DPB11/sld2-T84D::LEU2*<br>*bar1Δ::natNT2 PSF2-yeGFP::hphNT1* | this study |
| YKR1549 | *pep4Δ::kanMX4*<br>*his3::pGAL-DBF4::HIS3*<br>*leu2::pGAL-DPB11/sld2-T84D::LEU2* | this study |
| YKR1615 | *pep4Δ::kanMX4*<br>*his3::pGAL-DBF4::HIS3*<br>*leu2::pGAL-DPB11/sld2-T84D::LEU2*<br>*PSF2-yeGFP::hphNT1* | this study |
| YKR1815 | *leu2::pGAL-DPB11/sld2-T84D::LEU2*<br>*bar1Δ::natNT2*<br>*his3::pGAL-DBF4/SLD3::HIS3*<br>*trp1::pGAL-CDC45/SLD7::TRP1* | this study |
| YKR2113 | *leu2::pGAL-DPB11/sld2-T84D::LEU2*<br>*bar1Δ::natNT2*<br>*his3::pGAL-DBF4::HIS3*<br>*trp1::pGAL-JET1::TRP1* | this study |
| YKR2114 | *leu2::pGAL-DPB11/sld2-T84D::LEU2*<br>*bar1Δ::natNT2*<br>*his3::pGAL-DBF4/SLD3::HIS3*<br>*trp1::pGAL-JET1/SLD7::TRP1* | this study |
| YKR2035 | *leu2::pGAL-DPB11/sld2-T84D::LEU2*<br>*bar1Δ::natNT2*<br>*his3::pGAL-DBF4/SPT21-3FLAG::HIS3* | this study |
| YKR2090 | *his3::pGAL-DBF4::HIS3*<br>*leu2::pGAL-DPB11/sld2-T84D::LEU2*<br>*bar1Δ::natNT2*<br>*cdc45::JET1::TRP1* | this study |
| YKR2108 | *leu2::pGAL-DPB11/sld2-T84D::LEU2*<br>*bar1Δ::natNT2*<br>*cdc45::JET1::TRP1*<br>*his3::pGAL-DBF4/SPT21-3FLAG::HIS3* | this study |
| YKR1603 | *leu2::pGAL-DPB11/sld2-T84D::LEU2*<br>*bar1Δ::natNT2*<br>*his3::pGAL-dbf4-RxxL-4A (R10A,L13A,R62A,L65A)::HIS3* | this study |
| YKR1563 | *his3::pGAL-DBF4::HIS3*<br>*leu2::pGAL-DPB11/sld2-T84D::LEU2*<br>*bar1Δ::natNT2*<br>*rif1Δ::hphNT1* | this study |
| YKR1803 | *his3::pGAL-DBF4::HIS3*<br>*leu2::pGAL-DPB11/sld2-T84D::LEU2*<br>*bar1Δ::natNT2*<br>*sml1Δ::hphNT1*<br>*dif1Δ::kanMX4* | this study |
| YKR1614 | *leu2::pGAL-DPB11/sld2-T84D::LEU2*<br>*his3::pGAL-DBF4/RNR1::HIS3*<br>*bar1Δ::natNT2* | this study |
| YKR1824 | *leu2::pGAL-DPB11/sld2-T84D::LEU2*<br>*bar1Δ::natNT2*<br>*his3::pGAL-DBF4/rnr1-D57N::HIS3* | this study |
| YKR2099 | *his3::pGAL-DBF4::HIS3*<br>*leu2::pGAL-DPB11/sld2-T84D::LEU2*<br>*bar1Δ::natNT2*<br>*trp1::pGAL-CDC6::TRP1* | this study |
| YKR2059 | *trp1::pTDH3-TIR1-9myc,tTA,tetR'-SSN6::TRP1*<br>*SLD3::(hphNT1)iAID-promoter-sld3-3aid\*-9myc::natNT2* | this study |
| YKR2067 | *leu2::pGAL-DPB11/sld2-T84D::LEU2*<br>*his3::pGAL-DBF4::HIS3*<br>*trp1::pTDH3-TIR1-9myc,tTA,tetR'-SSN6::TRP1*<br>*SLD3::(hphNT1)iAID-promoter-sld3-3aid\*-9myc::natNT2* | this study |
| YKR1538 | *trp1::DDC1-RAD9-3FLAG::TRP1* | this study |
| YKR1541 | *his3::pGAL-DBF4::HIS3*<br>*leu2::pGAL-DPB11/sld2-T84D::LEU2*<br>*trp1::DDC1-RAD9-3FLAG::TRP1* | this study |
| YKR1564 | *trp1::DDC1-RAD9-3FLAG::TRP1*<br>*bar1Δ::natNT2* | this study |
| YKR1567 | *his3::pGAL-DBF4::HIS3*<br>*leu2::pGAL-DPB11/sld2-T84D::LEU2*<br>*trp1::DDC1-RAD9-3FLAG::TRP1*<br>*bar1Δ::natNT2* | this study |
| YKR2025 | *rad9Δ::hphNT1* | this study |
| YKR2026 | *mrc1Δ::hphNT1* | this study |
| YKR1484 | *his3::pGAL-DBF4::HIS3*<br>*leu2::pGAL-DPB11/sld2-T84D::LEU2*<br>*rad9Δ::hphNT1* | this study |
| YKR1481 | *his3::pGAL-DBF4::HIS3*<br>*leu2::pGAL-DPB11/sld2-T84D::LEU2*<br>*mrc1Δ::hphNT1* | this study |
| YKR1996 | *trp1::pGPD-TIR1-3myc::TRP1*<br>*cdc6-3aid\*-9myc::natNT2* | this study |
| YKR2000 | *trp1::pGPD-TIR1-3myc::TRP1*<br>*leu2::pGAL-DPB11/sld2-T84D::LEU2*<br>*his3::pGAL-DBF4::HIS3*<br>*cdc6-3aid\*-9myc::natNT2* | this study |
| YKR1754 | *ARS702::[I-SceI-cut site]::TRP1* | this study |
| YKR1755 | *leu2::pGAL-DPB11/sld2-T84D::LEU2*<br>*ARS702::[I-SceI-cut site]::TRP1* | this study |
| YKR1756 | *his3::pGAL-DBF4::HIS3*<br>*leu2::pGAL-DPB11/sld2-T84D::LEU2*<br>*ARS702::[I-SceI-cut site]::TRP1* | this study |
| YKR1642 | *rad52Δ::hphNT1* | this study |
| YKR1645 | *his3::pGAL-DBF4::HIS3*<br>*leu2::pGAL-DPB11/sld2-T84D::LEU2*<br>*rad52Δ::hphNT1* | this study |
| YKR1737 | *leu2::pADH-DPB11-VN::LEU2* | this study |
| YKR1768 | *leu2::pADH-DPB11-VN::LEU2*<br>*trp1::pADH-VC-sld2-T84A::TRP1* | this study |
| YKR1832 | *leu2::pADH-DPB11-VN::LEU2*<br>*trp1::pADH-sld2-T84A-VC::TRP1* | this study |
| YKR1767 | *leu2::pADH-DPB11-VN::LEU2*<br>*trp1::pADH-VC-SLD2::TRP1* | this study |
| YKR1831 | *leu2::pADH-DPB11-VN::LEU2*<br>*trp1::pADH-SLD2-VC::TRP1* | this study |
| YKR1830 | *leu2::pADH-DPB11-VC::LEU2* | this study |
| YKR1836 | *leu2::pADH-DPB11-VC::LEU2*<br>*trp1::pADH-VN-sld2-T84A::TRP1* | this study |
| YKR1834 | *leu2::pADH-DPB11-VC::LEU2*<br>*trp1::pADH-sld2-T84A-VN::TRP1* | this study |
| YKR1835 | *leu2::pADH-DPB11-VC::LEU2*<br>*trp1::pADH-VN-SLD2::TRP1* | this study |
| YKR1833 | *leu2::pADH-DPB11-VC::LEU2*<br>*trp1::pADH-SLD2-VN::TRP1* | this study |
| YKR1763 | W303-1A *CAN1::URA3*<br>*leu2::pADH-DPB11-VN::LEU2* | this study |
| YKR1783 | W303-1A *CAN1::URA3*<br>*leu2::pADH-DPB11-VN::LEU2*<br>*trp1::pADH-SLD2::TRP1* | this study |
| YKR1780 | W303-1A *CAN1::URA3*<br>*leu2::pADH-DPB11-VN::LEU2*<br>*trp1::pADH-VC-SLD2::TRP1* | this study |
| YKR1781 | W303-1A *CAN1::URA3*<br>*leu2::pADH-DPB11-VN::LEU2*<br>*trp1::pADH-VC-sld2-T84A::TRP1* | this study |
| YKR1850 | W303-1A *CAN1::URA3*<br>*leu2::pADH-DPB11-VN::LEU2*<br>*trp1::pADH-SLD2-VC::TRP1* | this study |
| YKR1851 | W303-1A *CAN1::URA3*<br>*leu2::pADH-DPB11-VN::LEU2*<br>*trp1::pADH-sld2-T84A-VC::TRP1* | this study |

### Plasmids

Genes of interest were amplified from genomic DNA of W303-1A and cloned into the respective vector using the In-Fusion HD cloning kit (Clontech). Mutations and deletions were introduced by oligonucleotide-directed site-specific mutagenesis.

**Table S2:** Plasmids used in this study.

| **plasmid name** | **vector** | **insert** |
| --- | --- | --- |
| pKR588 | pFA6a-natNT2 | *3aid\*-9myc* |
| pKR534 | YIplac128 | *pGAL-DPB11/SLD2* |
| pKR535 | YIplac128 | *pGAL-DPB11/sld2-T84D* |
| pKR520 | pRS303 | *pGAL-DBF4* |
| pKR546 | pRS303 | *pGAL-DBF4/SLD3* |
| pKR581 | YIplac204 | *pGAL-CDC45/SLD7* |
| pKR609 | YIplac204 | *pGAL-JET1/SLD7* |
| pKR608 | YIplac204 | *pGAL-JET1* |
| pKR598 | pRS303 | *pGAL-SPT21-3FLAG* |
| pKR592 | pRS303 | *pGAL-DBF4/SPT21-3FLAG* |
| pKR604 | YIplac204 | *pGAL-PRI1/PRI2* |
| pKR614 | pRS303 | *pGAL-POL1-3FLAG/POL12* |
| pKR531 | pRS303 | *pGAL-dbf4ΔD-box* |
| pKR563 | pRS303 | *pGAL-RNR1* |
| pKR582 | pRS303 | *pGAL-rnr1-D57N* |
| pKR562 | pRS303 | *pGAL-DBF4/RNR1* |
| pKR583 | pRS303 | *pGAL-DBF4/rnr1-D57N* |
| pKR508 | YIplac204 | *pGAL-CDC6* |
| pKR548 | pRS304 | *pGPD-TIR1-3myc* |
| pKR385 | YIplac128 | *pADH-DPB11-VN* |
| pKR386 | YIplac128 | *pADH-DPB11-VC* |
| pKR477 | YIplac204 | *pADH-SLD2* |
| pKR417 | YIplac204 | *pADH-VC-SLD2* |
| pKR425 | YIplac204 | *pADH-VC-sld2-T84A* |
| pKR403 | YIplac204 | *pADH-SLD2-VC* |
| pKR413 | YIplac204 | *pADH-sld2-T84A-VC* |
| pKR416 | YIplac204 | *pADH-VN-SLD2* |
| pKR424 | YIplac204 | *pADH-VN-sld2-T84A* |
| pKR402 | YIplac204 | *pADH-SLD2-VN* |
| pKR412 | YIplac204 | *pADH-sld2-T84A-VN* |

### Flow cytometry

About 10^7^ cells (0.5–1 OD) were harvested by centrifugation, resuspended in 50 mM Tris-HCl pH 8.0/70% ethanol and stored at 4 °C for at least one hour for fixation and permeabilization. Afterwards, cells were digested with RNAseA buffer (50 mM Tris-HCl pH 8.0, 0.38 mM MgCl_2_, 0.38 mg/ml RNase A (Sigma R4875)) overnight at 37 °C and with proteinase K buffer (50 mM Tris-HCl pH 8.0, 5% glycerol, 2.5 mM CaCl_2_, 1 mg/ml proteinase K (Sigma P2308)) for 30 min at 50 °C. Cells were resuspended in 50 mM Tris-HCl pH 8.0, sonicated, diluted 1:20 with 50 mM Tris-HCl pH 8.0 containing 0.5 µM SYTOX green (Invitrogen S7020), and measured on a MACSquant analyzer (Miltenyi Biotec).

To measure DNA synthesis via flow cytometry, EdU-treated cells were processed analogously to samples for cell cycle analysis and afterwards incubated for 60 min in PBS supplemented with 1% BSA. One half was subjected to a click chemistry reaction with disulfo-Cy5-picolyl-azide (Jena Bioscience CLK-1177) for one hour, whereas the other half was kept as a control. A click chemistry reaction for 10^7^ cells (1 OD) consisted of 36 µl PBS, 2 µl freshly prepared 1 M ascorbic acid, 2 µl 1 M CuSO_4_ and 0.5 µl 2 mM disulfo-Cy5-picolyl-azide. After the click chemistry reaction, the cells were washed twice with 10% ethanol in PBS before they were resuspended in PBS. Both the click chemistry reaction and the control sample were diluted 1:20 with SYTOX buffer (50 mM Tris-HCl pH 8.0, 0.5 µM SYTOX green) and measured on a MACSquant analyzer (Miltenyi Biotec).

Flow cytometry data were analyzed and plotted using FlowJo (v10.6.2). For quantification, B1 channel (SYTOX green fluorescence) measurements were exported and fitted to a bimodal distribution model with one population anchored on the 1C DNA content peak using the mixtools package (v1.2.0) (Benaglia et al., 2009) in R (v4.0.3).

### EdU-IP for sequencing

For each sample, approximately 10^9^ cells (100 OD) were harvested by centrifugation, fixed with 50 mM Tris-HCl pH 8.0/70% ethanol for at least one hour, and then digested with 25 ml RNaseA buffer (50 mM Tris-HCl pH 8.0, 0.38 mM MgCl_2_, 0.38 mg/ml RNase A) overnight at 37 °C. Afterwards, cells were washed with 50 mM Tris-HCl pH 8.0, digested with 10 ml proteinase K buffer (50 mM Tris-HCl pH 8.0, 5% glycerol, 2.5 mM CaCl_2_, 1 mg/ml proteinase K) for one hour at 50 °C, and subsequently incubated with 25 ml PBS supplemented with 1% BSA for another hour at room temperature. The cells were afterwards subjected to an upscaled click chemistry reaction with biotin-picolyl-azide (Jena Bioscience CLK-1167) for one hour at room temperature and washed twice with 10% ethanol in PBS afterwards. Next, cells were resuspended in breaking buffer (2% triton X-100, 1% SDS, 100 mM NaCl, 10 mM Tris-HCl pH 8.0, 1 mM EDTA) and subjected to mechanical lysis. DNA from this lysate was sheared to 300 bp fragments using a BioRuptor UCD-200 sonicator (Diagenode). Cell debris were removed by high-speed centrifugation and DNA from the supernatant was isolated by ethanol precipitation and resuspension in TE buffer. Labeling of the DNA with biotin was confirmed in dot blots using HRP-conjugated streptavidin (Sigma S5512, 1 µg/ml) for detection. The size distribution of DNA fragments was analyzed by agarose gel electrophoresis and 20 µl were taken aside for library preparation (total DNA).

Equal amounts (approx. 700 ng) of sheared, EdU-biotin-labeled DNA were mixed 1:1 with 2x WB buffer (10 mM Tris-HCl pH 8.0, 10 mM EDTA, 1 M NaCl, 0.02% NP-40) supplemented with 1 mg/ml BSA and incubated with 25 µl of streptavidin-coupled magnetic beads (Thermo Fisher Scientific Dynabeads M-280) for 30 min at room temperature. The beads were washed five times for 5 min with 1x WB buffer (diluted with TE buffer). Subsequently, the beads were eluted twice with 100 µl buffer EB (10 mM Tris-HCl pH 8.0, 10 mM EDTA, 0.1% SDS) for one hour at 55 °C. The eluates were pooled, purified by phenol/chloroform/isoamyl alcohol extraction and precipitated in the presence of 50 µg/ml GlycoBlue coprecipitant (Invitrogen AM9515) with sodium acetate and absolute ethanol. After drying, the pellet was resuspended in 20 µl TE buffer.

Libraries for next-generation sequencing were prepared using the NEBNext Ultra II DNA library prep kit (New England Biolabs) following the manufacturer’s instructions and sequenced on an Illumina NextSeq 500 platform (75 bp reads, paired end) at the MPIB NGS core facility.

### Western blots

Approximately 2×10^7^ cells (1 OD) were harvested by centrifugation and snap-frozen in liquid nitrogen. Afterwards, cells were resuspended in 1 ml water, supplemented with 150 µl 1.85 M NaOH and 7.5% β-mercaptoethanol and incubated for 15 min at 4 °C. Subsequently, 150 µl 55% tri-chloroacetic acid (TCA) were added for 10 min at 4 °C, before collecting the pellet, resuspending it in 50 µl HU buffer (8 M urea, 5% SDS, 200 mM Tris-HCl pH 6.8, 1.5% DTT, bromophenolblue), and heating it for 10 min at 65 °C.

Samples were loaded on NuPAGE 4-12% Bis-Tris acrylamide gels (Invitrogen NP0322) and run at 200 V with MOPS buffer or MES buffer, according to the proteins that needed to be separated. To resolve phosphorylated isoforms of Rad53, standard 10% acrylamide gels were run with SDS buffer.

After gel electrophoresis, proteins were transferred to nitrocellulose membranes using a tank blot system and methanol-containing transfer buffer. The transfer was carried out at 4 °C with 90 V for 90 min. After transfer, primary antibodies were diluted in superblotto (2.5% skim milk powder, 0.5% BSA, 0.5% NP-40, 0.1% tween-20 in TBS) and added to the membranes for incubation overnight at 4 °C. After washing once for 5 min with western wash buffer, secondary antibodies (diluted 1:3000 in superblotto) were added for 90 min at room temperature. For detection of the immune-blots, Pierce ECL western blotting substrate (Thermo Fisher Scientific 32106) was added following the manufacturer’s instructions and chemiluminescence was detected using a LAS-3000 CCD camera system (Fujifilm).

Primary antibodies used in this study are: anti-γH2A (abcam ab181447, rabbit, 1:2000 dilution), anti-Rad53 (abcam ab104232, rabbit, 1:4000 dilution), anti-Dpb11 (BPF19, (Pfander and Diffley, 2011)), rabbit, 1:5000 dilution), anti-Sld2 (kind gift of Philip Zegerman, rabbit, 1:2000 dilution).

### Immunoprecipitation of replisomes

Experiments were done as triplicates and, for each sample, 2×10^9^ cells (100 OD per sample) were stopped by the addition of 0.1% NaN_3_ and kept on ice for 30 min before being harvested by centrifugation. Replisomes were purified based on previously published work (Gambus et al., 2006). Briefly, cells were washed with 10 mM HEPES-KOH pH 7.9, resuspended in lysis buffer (100 mM HEPES-KOH pH 7.9, 50 mM potassium acetate, 10 mM magnesium acetate, 2 mM EDTA) including protease inhibitors, and snap-frozen as yeast popcorn in liquid nitrogen. The yeast popcorn was ground to fine powder using a cryogenic mill (SPEX SamplePrep), thawed and supplemented with 0.25 volumes glycerol mix buffer (100 mM HEPES-KOH pH 7.9, 300 mM potassium acetate, 10 mM magnesium acetate, 2 mM EDTA, 50% glycerol, 0.5% NP-40) to obtain an extract with 10% glycerol, 100 mM potassium acetate and 0.1% NP-40. After incubation with 800 U/ml SmDNase (MPIB core facility) for 30 min on ice, the extract was cleared by centrifugation. The protein concentration was measured using a standard Bradford assay and, after adjusting the concentrations, the extracts were used directly for immunoprecipitation.

Agarose GFP-trap beads (Chromotek gta-100, 20 µl used per sample) were equilibrated with IP wash buffer (100 mM HEPES-KOH pH 7.9, 100 mM potassium acetate, 10 mM magnesium acetate, 2 mM EDTA, 10% glycerol) including 0.1% NP-40 and then incubated with 30 mg of total protein for two hours at 4 °C. Afterwards, the beads were washed three times with IP wash buffer including NP-40 and two times with IP wash buffer lacking NP-40.

### Mass spectrometry measurement

Washed beads were incubated for 30 min with elution buffer 1 (2 M urea, 50 mM Tris-HCl pH 7.5, 2 mM DTT, 20 µg/ml trypsin) followed by a second elution with elution buffer 2 (2 M urea, 50 mM Tris-HCl pH 7.5, 10 mM chloroacetamide) for 5 min. Both eluates were combined and further incubated at room temperature overnight. Tryptic peptide mixtures were acidified to 1% TFA and desalted with Stage Tips containing C18 reverse-phase material and analyzed by mass spectrometry.

Peptides were separated on 50 cm columns packed with ReproSil-Pur C18-AQ 1.9 μm resin (Dr. Maisch GmbH). Liquid chromatography was performed on an EASY-nLC 1200 ultra-high-pressure system coupled through a nano-electrospray source to a Q-Exactive HF-X Mass Spectrometer (Thermo Fisher Scientific). Peptides were loaded in buffer A (0.1% formic acid) and separated with a non-linear gradient of 5-60 % buffer B (0.1% formic acid, 80% acetonitrile) at a flow rate of 300 nl/min over 50 min. The column temperature was kept at 60 °C by an in-house designed oven with a Peltier element. Data acquisition switched between a full scan (60 K resolution, 20 ms max. injection time, AGC target 3e6) and 10 data-dependent MS/MS scans (15K resolution, 60 ms max. injection time, AGC target 1e5). The isolation window was set to 1.4 and normalized collision energy to 27. Multiple sequencing of peptides was minimized by excluding the selected peptide candidates for 30 s.

### Pulsed-field gel electrophoresis and Southern blotting

Pulsed-field gel electrophoresis and Southern blotting was performed with modifications as in (Bittmann et al., 2020). For each timepoint, approximately 4×10^7^ cells (2 OD) were harvested, resuspended in ice-cold Stop Buffer (150 mM NaCl, 50 mM NaF, 2 mM NaN_3_, 10 mM EDTA) and stored at 4 °C until further processing. Samples were washed twice with ice-cold 50 mM EDTA, resuspended in SCE buffer (1 M sorbitol, 0.1 M sodium citrate, 10 mM EDTA) + 150 U/ml zymolyase 100T (Roth 9329) and mixed with 50 µl 2% agarose before casting into plugs. After solidification, plugs were placed in SCEM (1 M sorbitol, 0.1 M sodium citrate, 10 mM EDTA, 5% β-mercaptoethanol) + 150 U/ml zymolyase 100T and incubated at 37 °C for 2 days. Afterwards, plugs were washed with TE (10 mM Tris-HCl pH 8.0, 1 mM EDTA) for 1-2 h each wash, placed into PK buffer (1 mg/ml sarcosyl, 0.5 M EDTA, 2 mg/ml proteinase K) and incubated at 55 °C for 2 days. Plugs were washed three more times with TE before use.

Plugs were loaded on a gel containing 1% agarose (Bio-Rad Cat. 1620138) in 0.5x TBE (45 mM Tris, 45 mM borate, 0.5 mM EDTA). Electrophoresis was carried out in 14 °C cold 0.5x TBE in a CHEF DR-III system (Bio-Rad, initial switch time 60 sec, final switch time 120 sec, 6 V/cm, angle 120°, 24 h). Afterwards, the gel was stained with 1 μg/ml ethidium-bromide in 0.5x TBE for one hour and de-stained with deionized water. Images were taken using a GenoSmart gel documentation system (VWR).

For Southern blotting, the DNA was nicked in 0.125 M HCl for 10 min, denatured in 1.5 M NaCl, 0.5 M NaOH for 30 min and neutralized by 0.5 M Tris, 1.5 M NaCl (pH 7.5) for 30 min. The DNA was transferred onto a Hybond-N+ membrane (GE healthcare) and UV-cross-linked (Stratagene Stratalinker 1800, auto-crosslink function). The membrane was probed with a radioactive (α-^32^P dCTP) labeled *TRP1* fragment and imaged using a Typhoon FLA 9000 imaging system (GE Healthcare).

### Strand-specific RPA-ChIP-seq

Samples for strand-specific RPA-ChIP-seq were prepared as described previously (Peritore et al., 2021). Briefly, 2×10^9^ cells (100 OD) were crosslinked at the indicated timepoints with 1% formaldehyde for 16 min at room temperature, subsequently quenched with 400 mM glycine for 60 min, washed with PBS and frozen in liquid nitrogen. After resuspending in lysis buffer (50 mM HEPES-KOH pH 7.5, 150 mM NaCl, 1 mM EDTA, 1% triton X-100, 0.1% sodium-deoxycholate, 0.1% SDS), cells were mechanically lysed and chromatin was sheared to 200-500 bp fragments. Cell lysates were cleared by centrifugation and diluted 1:1 with lysis buffer. 1% of the extract was taken as a total DNA sample and 40% of the extract were incubated with an antibody against budding yeast RFA (Agrisera, AS07 214) for 2 hours followed by 30 minutes incubation with protein A-coupled dynabeads (Invitrogen 10002D). Beads were washed three times with lysis buffer, once with lysis buffer supplemented with 500 mM NaCl, once with wash buffer (10 mM Tris-HCl pH 8.0, 0.25 M LiCl, 1 mM EDTA, 0.5% NP-40, 0.5% sodium-deoxycholate), and once with TE pH 8.0. Immunoprecipitated complexes were eluted with 1% SDS, proteins were degraded with proteinase K and crosslinks were reversed at 65 °C. DNA was purified by phenol-chloroform extraction and cleaned up using Phase Lock Gel tubes (5Prime) and ethanol precipitation.

Strand-specific ChIP-seq libraries were prepared from 1-3 ng of DNA using Accel-NGS 1S Plus Library Kit (Swift Biosciences) following the manufacturer’s instructions and sequenced on an Illumina NextSeq 500 (75 bp or 37 bp reads, paired-end) at the MPIB NGS core facility.

### Split-Venus fluorescence intensity measurement and quantification

Cells were grown at 30 °C in YPD supplemented with adenine to log-phase (OD600 of 0.5-0.6), stopped by adding 0.1% NaN_3_, and kept in the dark on ice for 30 min. After two washes with 50 mM Tris-HCl pH 8.0, cells were re-suspended in 50 mM Tris-HCl pH 8.0 and measured on a MACSquant analyzer (Miltenyi Biotech). Values for mean as well as 97^th^ percentile fluorescence intensity were calculated and exported using FlowJo (v10.6.2) and data from 6 independent cultures per strain were used to generate boxplots using R (v4.0.3) after subtracting background fluorescence as measured in a strain that expressed an untagged *SLD2* construct.

### Gross chromosomal rearrangement assay and rate calculation

Rates of gross chromosomal rearrangements (GCRs) were determined using a standard protocol (Putnam and Kolodner, 2010). Briefly, pre-cultures of *S. cerevisiae* cells harboring a *CAN1::URA3* reporter on chromosome 5 were grown in SC-Ura medium and plated out on YPD plates to obtain colonies that formed from single cells. Eight colonies were excised from the plates for each condition and used to inoculate larger cultures in YPD (control strains: 50 ml; strains with stabilized interaction: 2 ml), which were grown to stationary phase at 30 °C. The number of viable cells was determined by plating a serial dilution (10^-6^) on non-selective YPD plates. The total number of GCR events was determined by plating the remaining culture on SC-Arg plates that were supplemented with 50 mg/L L-canavanine (Sigma C9758) and 1 g/L 5’-fluoroorotic acid (US Biological Life Sciences F5050) to select against both *CAN1* and *URA3*. No more than 10^9^ cells were spread on each selection plate and the plates were incubated at 30 °C for two days (YPD) and three to five days (selection). Afterwards, the clones were counted and GCR rates as well as confidence intervals were calculated by fluctuation analysis using the maximum likelihood method in the web tool FALCOR (B. M. Hall et al., 2009) that was kindly made accessible by the Liang lab under https://lianglab.brocku.ca/FALCOR/.

### Mass spectrometry data analysis

Raw mass spectrometry data were analyzed with MaxQuant (v1.5.3.54) (Cox and Mann, 2008). Peak lists were searched against the yeast Uniprot FASTA database combined with 262 common contaminants by the integrated Andromeda search engine. The false discovery rate was set to 1% for both peptides (minimum length of 7 amino acids) and proteins. “Match between runs” (MBR) with a maximum matching time window of 0.5 min and an alignment time window of 20 min was enabled. Relative protein amounts were calculated with the MaxLFQ algorithm with a minimum ratio count of two.

Absolute protein intensity (iBaq) estimates were calculated dividing the LFQ intensities by the theoretical number of tryptic peptides of each protein.

Statistical analysis of LFQ-derived protein expression data was performed using R. LFQ values were log2 transformed. Per each experimental condition, a pairwise comparison was performed with triplicate pulldowns of bait (GFP-Psf2) vs. control (untagged Psf2) yeast strains. For each comparison, the dataset was filtered to present at least two valid values in the bait group and missing values imputed with a downshift of 1.8 standard deviations and a width of 0.2 standard deviations. Fold changes and p-values were calculated using an unpaired Student’s T-Test.

### Next-generation sequencing data analysis

For each sample about 10 million sequencing reads were obtained and quality-checked using FastQC (v0.11.9, https://www.bioinformatics.babraham.ac.uk/projects/fastqc/). The reads were aligned to the budding yeast reference genome sacCer3 (Engel et al., 2014) using the Burrows-Wheeler aligner bwa (v0.7.17) with standard parameters (H. Li and Durbin, 2010) and the alignments were sorted and indexed using samtools (v1.12) (H. Li et al., 2009). The tool bamCoverage from the deepTools suite (v3.5.1) (Ramírez et al., 2016) was used with the options “--binSize 50 --minMappingQuality 60 --normalizeUsing CPM” to calculate coverage normalized to sequencing depth per 50 bp bins and discard multi-mapping reads. Reads mapping to the rDNA locus were blacklisted during this step. The tool bigwigCompare was used afterwards to normalize samples to total input DNA. Locations of origins of replication were used as annotated in oriDB (Siow et al., 2011) and annotations for telomeres and centromeres were taken from SGD (Cherry et al., 2012). Data were plotted using plotHeatmap from the deepTools suite or pyGenomeTracks (v3.6) (Lopez-Delisle et al., 2021).

To separate reads by strands, alignment files were filtered with samtools using the options “-f 99” and “-f 147” for reads mapping to the forward strand and the options “-f 83” and “-f 163” for reads mapping to the reverse strand before calculating bigWig-coverage files. To calculate asymmetry profiles, the log2 ratio of depth-normalized forward and reverse reads was first calculated and then averaged over all chromosomes using computeMatrix and plotHeatmap/plotProfile from deepTools.

To calculate a normalized asymmetry score per chromosome, the average absolute value of asymmetry (log2(forward/reverse)) per 50 bp bin was calculated and plotted against total chromosome length using R (v4.0.3).

### Data availability

All sequencing data have been deposited as raw fastq-files as well as depth-normalized bigwig-files and are available via accession number GSE182203. Mass spectrometry data is available from EBI-PRIDE via accession number.

---

**Figure S1 (related to Figure 1).**
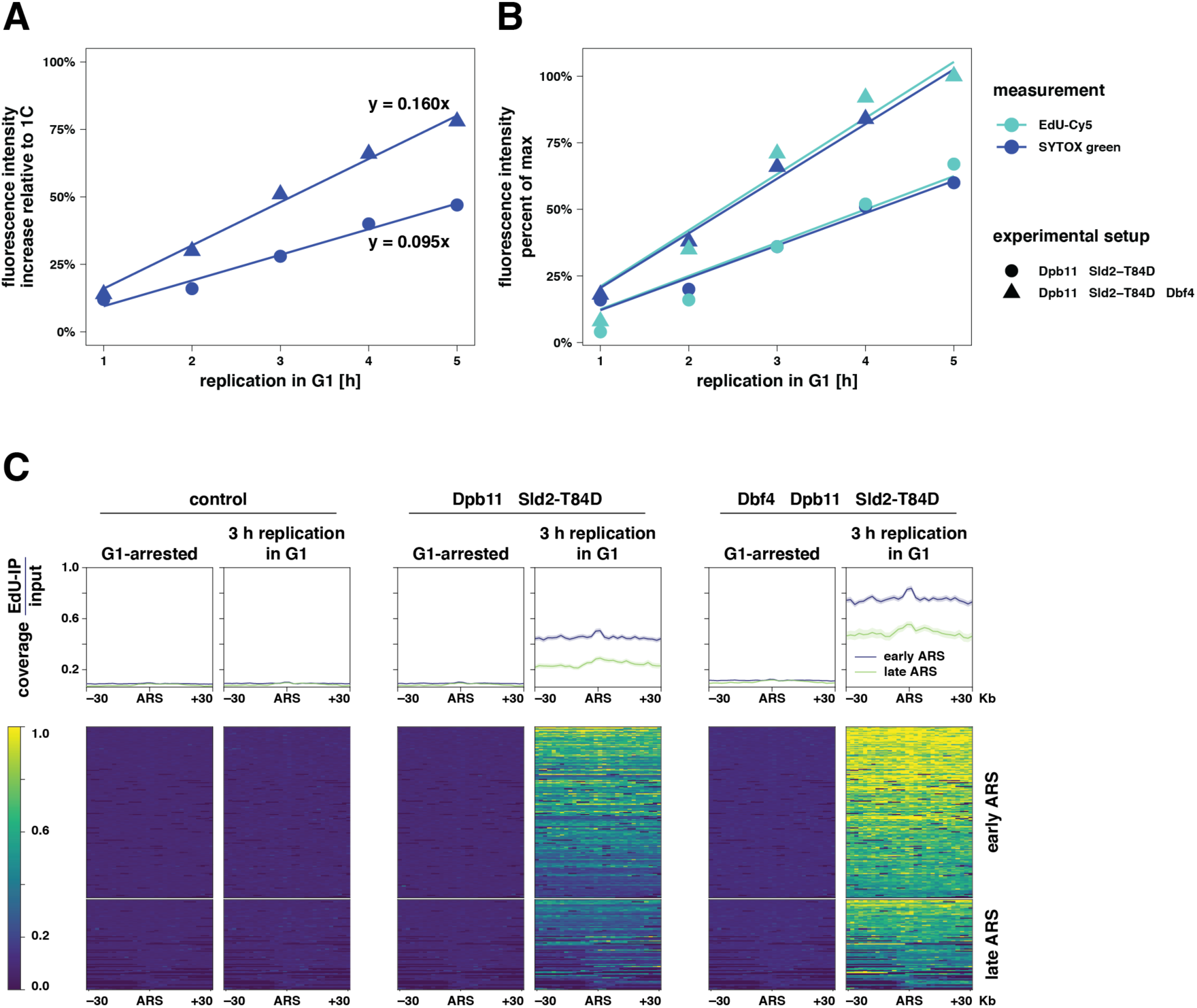
(A) DNA content increases linearly during unscheduled replication in G1. Related to Fig. 1B. Mean fluorescence intensity of total DNA (SYTOX green, blue) was measured by flow cytometry after induction of G1 replication for the indicated amount of time. Data were corrected for background mitochondrial DNA synthesis in a control strain, normalized to a DNA content of 1 C and fitted with a linear regression model (see equation). (B) DNA is quantitatively labeled with EdU during G1 replication. Related to Fig. 1B. As in (A), additionally measuring mean fluorescence intensity of newly synthesized DNA in G1 (EdU-Cy5, turquoise). Data were corrected for background mitochondrial DNA synthesis in a control strain and normalized to the respective maximum value. (C) Early-replicating origins are activated during G1 replication. Related to Fig. 1C. Input-normalized EdU-sequencing data were analyzed at early or late replication origins (ARS) ± 30 Kb. (top) Profile plots of mean coverage (dark) ± SE (light). (bottom) Heatmaps with 1 Kb bin size. Data from n=2 replicates.

**Figure S2 (related to Figure 2).**
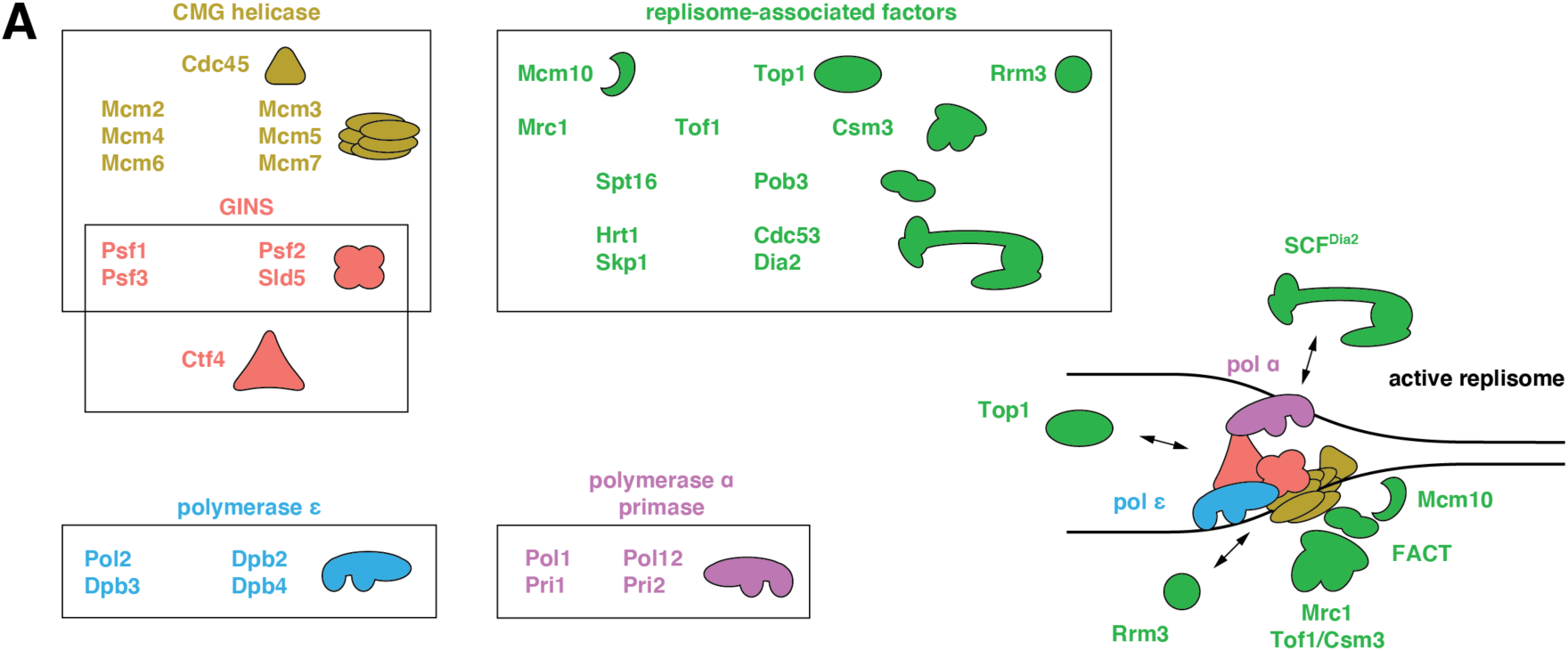
(A) Schematic drawing of protein complexes associating with GINS/Ctf4 in the context of a replisome. Color code as used in Fig. 2. Interaction of GINS/Ctf4 with Mcm2-7/Cdc45 indicates formation of the CMG helicase, which is the key regulated step in the transition from inactive helicase precursors to active replisomes during replication initiation. DNA polymerases and replisome-associated factors are recruited during this transition and travel with the replisome.

**Figure S3 (related to Figure 3).**
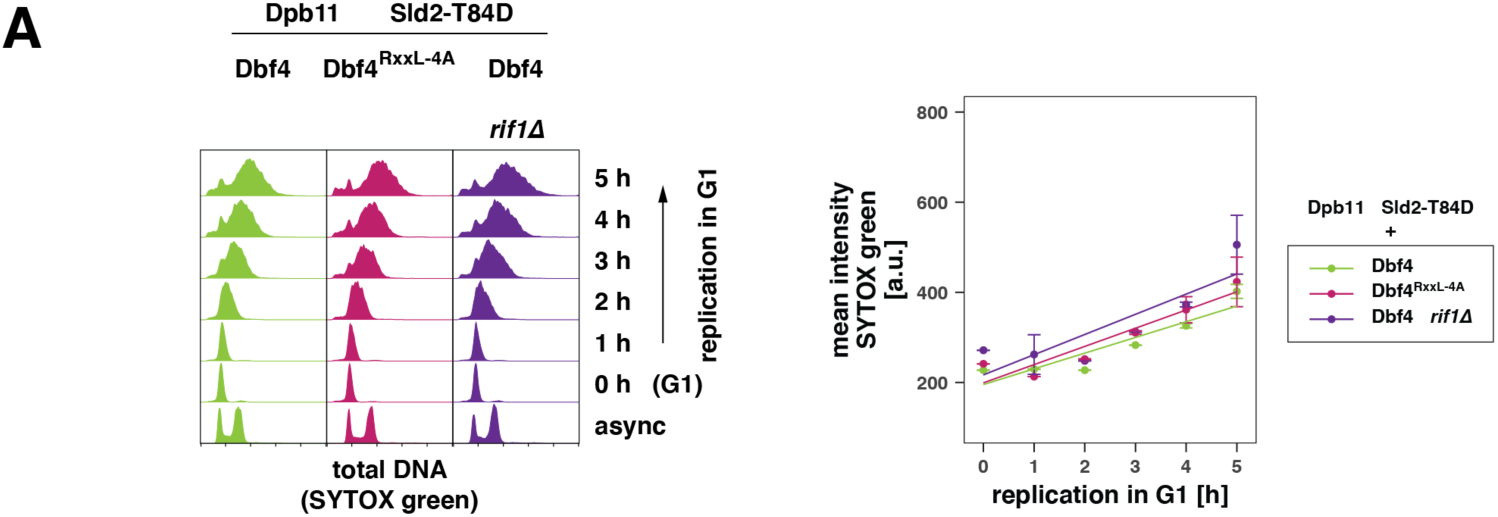
(A) Further deregulation of Dbf4-dependent kinase (DDK) does not increase the amount of unscheduled replication in G1 in the basic setup of CDK/DDK bypass. Experiment as in Fig. 1B but additionally using a strain expressing a *DBF4* allele with mutated destruction-boxes (*dbf4^RxxL-4A^*) or lacking the targeting subunit *RIF1* for the DDK-antagonizing PP1 phosphatase. (left) SYTOX green-stained total DNA after induction of replication by galactose-inducible promoter driven expression of the indicated proteins in G1-arrested cells as measured by flow cytometry at the indicated timepoints. (right) Quantification of the data by approximation of a bimodal distribution and calculating the means of the individual normal distributions. The average mean from 5 fits ± SD per timepoint is shown together with a linear regression.

**Figure S4 (related to Figure 4).**
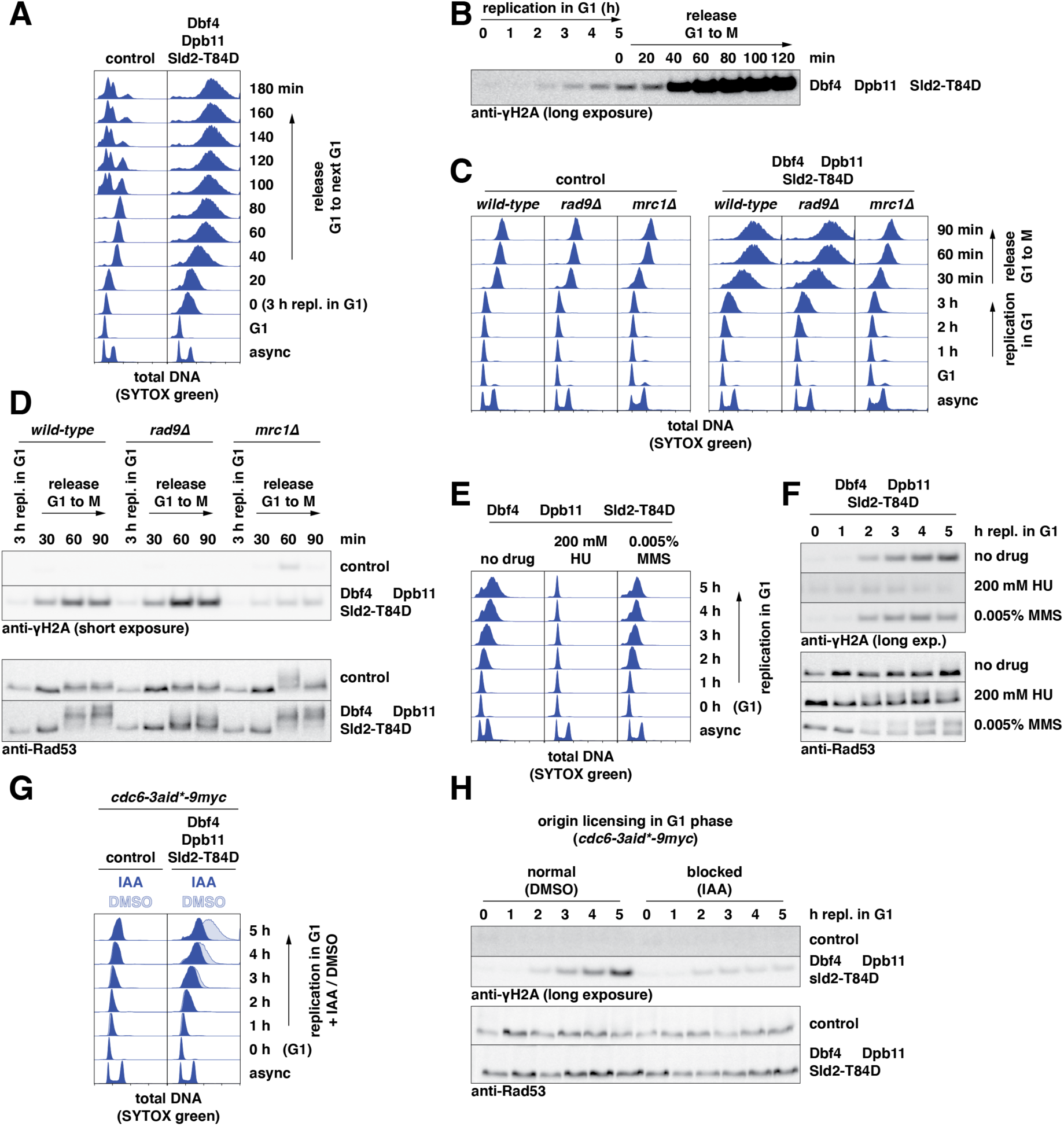
(A) Release from unscheduled G1 replication results in a cell cycle arrest after S phase. SYTOX green-stained total DNA after induction of replication in G1 and release to the next G1 as measured by flow cytometry at the indicated timepoints. Starting at 80 min control cells enter next G1-phase, while G1 replication cells stay arrested with G2/M DNA content. (B) Unscheduled G1 replication generates low amounts of DNA damage already in G1. Longer exposure of the γH2A western blot that is shown in Fig. 4B. (C) and (D) DNA damage after unscheduled replication in G1 is detected by the Rad9-dependent branch of the checkpoint. Unscheduled replication in G1 was induced by CDK/DDK bypass before release of the cells to M (nocodazole). (C) SYTOX green-stained total DNA content as measured by flow cytometry at the indicated timepoints. (D) Western blots of samples from (C) detecting γH2A and Rad53 at the indicated timepoints. (E) and (F) The checkpoint detects canonical replication stress in G1-arrested cells. (E) Cells undergoing unscheduled G1 replication were exposed to either hydroxyurea (HU) or methyl methanesulfonate (MMS) and total DNA was stained with SYTOX green and measured by flow cytometry. (F) Western blots of samples from (E) detecting γH2A and Rad53 at the indicated timepoints. (G) and (H) Generation of DNA damage in G1 requires continuous origin licensing. (G) SYTOX green-stained total DNA after induction of replication by galactose-inducible promoter driven expression of the indicated proteins in G1-arrested cells as measured by flow cytometry at the indicated timepoints. Cells were depleted of licensing factor Cdc6 at the beginning of the time course using an auxin-inducible degron allele. (H) Western blots of samples from (G) detecting γH2A and Rad53 at the indicated timepoints.

**Figure S5 (related to Figure 5).**
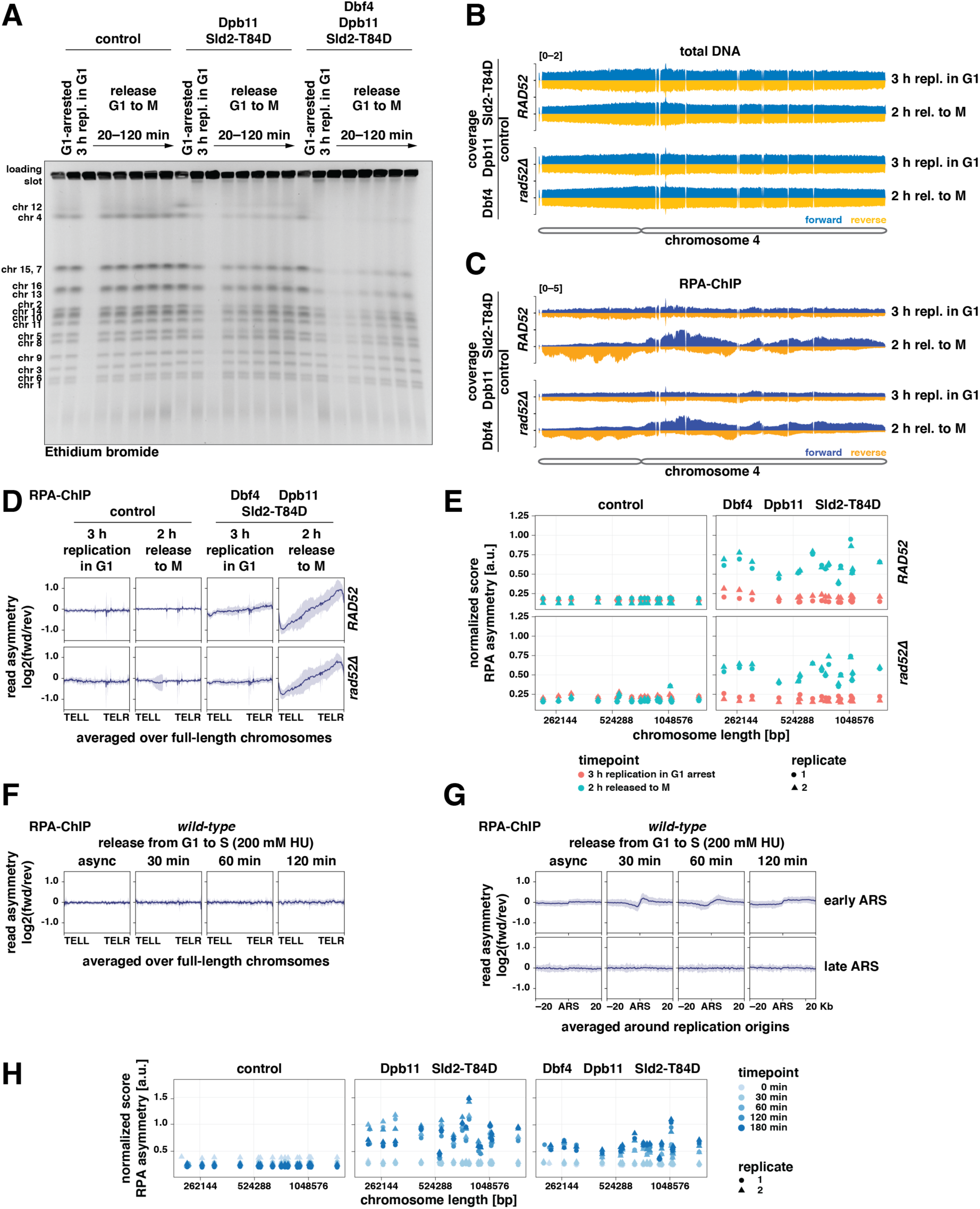
(A) Persistent replication/repair structures after release from unscheduled G1 replication. Ethidium bromide-stained gel corresponding to Fig. 5A. Samples were taken at the indicated timepoints after inducing G1 replication and releasing to M (nocodazole) and separated on a pulsed-field electrophoresis gel. (B) to (E) The strand-biased RPA pattern does not depend on *RAD52*. (B) and (C) show representative coverage traces of reads mapping to chromosome 4 for total DNA (B) and RPA-ChIP (C) of strains with and without *RAD52*. (D) The log2-ratio of RPA-ChIP-seq reads mapping to forward and reverse strands was averaged over full-length chromosomes at the indicated timepoints and is plotted as mean (dark) ± SD (light). (E) RPA asymmetry scores for each chromosome were calculated by normalizing the log2-ratios of RPA-ChIP-reads mapping to forward and reverse strand in 50 bp bins for chromosome length. Data from n=2 replicates. (F) and (G) Stalled replication forks cause only little strand-biased RPA binding to chromosomes around early-replicating origins. RPA-ChIP samples taken from *wild-type* cells after G1-arrest and release to S-phase in the presence of 200 mM HU for the indicated times. (F) Asymmetric RPA-ChIP read distribution to forward and reverse strands averaged over full-length chromosomes as in (D). (G) Asymmetric RPA-ChIP read distribution as in (D)/(F) but calculated around early- and late-replicating origins (autonomously replicating sequences (ARS)) ± 20 Kb. (H) Long and short chromosomes are equally affected by asymmetric RPA accumulation. RPA asymmetry scores for each chromosome were calculated by normalizing the log2-ratios of RPA-ChIP-reads mapping to forward and reverse strand in 50 bp bins for chromosome length. Data from n=2 replicates.

**Figure S6 (related to Figure 6).**
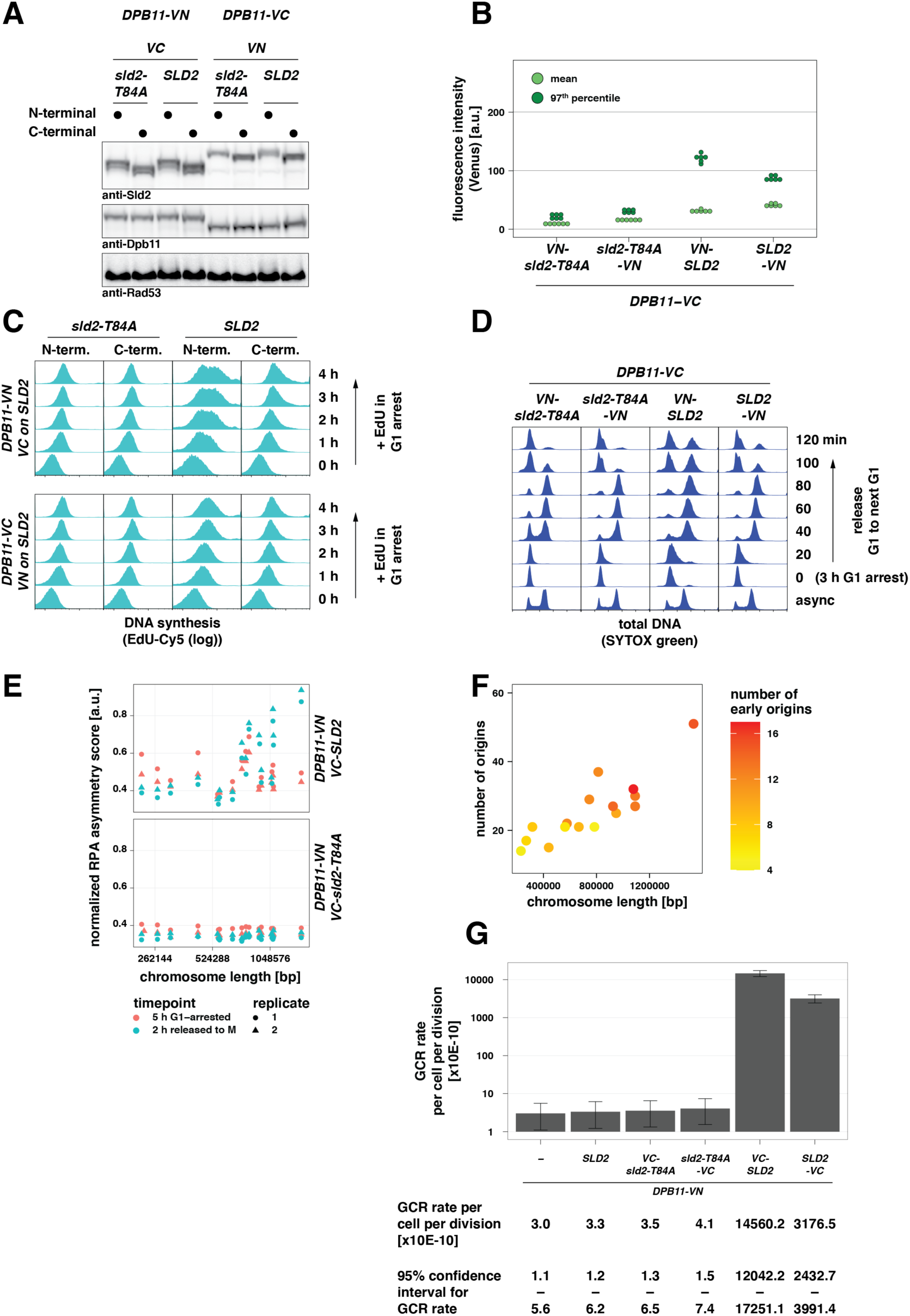
(A) Split-Venus tagged Dpb11- and Sld2-constructs are expressed to similar levels. Expression levels of Sld2 and Dpb11 carrying split-Venus tags as well as phosphorylated Rad53 detected by western blots from log-phase samples. Note the faint signal for phosphorylated Rad53 with constructs that stabilize the physical interaction between Dpb11 and Sld2. (B) Split-Venus tags (VN/VC) stabilize the physical interaction between Dpb11 and Sld2. Dpb11-VC and Sld2 tagged at either N- or C-terminus with VN fragment of the fluorescent protein Venus. Data represents the resulting mean (light green) and 97^th^ percentile (dark green) of split-Venus fluorescence intensity measured by flow cytometry in log-phase cells from n=6 replicates. (C) Venus-stabilized interaction of Dpb11 and Sld2 induces DNA replication in G1. Cells of the indicated genotypes were pre-arrested in G1 for 1 h and then kept arrested in G1 in the presence of EdU. Incorporated EdU was labeled with Cy5 and measured by flow cytometry. Note the logarithmic scaling of the x-axis to resolve small amounts of G1 replication. (D) Venus-stabilized interaction of Dpb11 and Sld2 results in cell cycle arrest. Additional samples to experiment shown in Fig 6B. SYTOX green-stained total DNA from samples at the indicated timepoints after release from G1 arrest to the next G1-phase as measured by flow cytometry. (E) Sporadic G1 replication affects long chromosomes more strongly. RPA asymmetry scores for each chromosome were calculated by normalizing the log2-ratios of RPA-ChIP-seq reads mapping to forward and reverse strand in 50 bp bins for chromosome length. Data from n=2 replicates. (F) Long chromosomes harbor more early-firing origins. Length of chromosomes was plotted against the total number of ARS sequences. Color intensity indicates the number of early-firing origins. (G) High levels of genome instability are caused by Venus-stabilized interaction of Dpb11 and Sld2. GCR rates for the assay shown in Fig. 6F were calculated from n=8 cultures by fluctuation analysis. Error bars indicate a 95% confidence interval for the determined GCR rate. Note the logarithmic scaling of the y-axis.

## References

Alexander, J.L., Barrasa, M.I., Orr-Weaver, T.L., 2015. Replication Fork Progression during Re-replication Requires the DNA Damage Checkpoint and Double-Strand Break Repair. Current Biology 25, 1654–1660. doi:10.1016/j.cub.2015.04.058

Alexander, J.L., Beagan, K., Orr-Weaver, T.L., McVey, M., 2016. Multiple mechanisms contribute to double-strand break repair at rereplication forks in Drosophila follicle cells. Proc. Natl. Acad. Sci. U.S.A. 113, 13809–13814. doi:10.1073/pnas.1617110113

Archambault, V., Ikui, A.E., Drapkin, B.J., Cross, F.R., 2005. Disruption of mechanisms that prevent rereplication triggers a DNA damage response. Mol. Cell. Biol. 25, 6707–6721. doi:10.1128/MCB.25.15.6707-6721.2005

Arias, E.E., Walter, J.C., 2005. Replication-dependent destruction of Cdt1 limits DNA replication to a single round per cell cycle in Xenopus egg extracts. Genes Dev. 19, 114–126. doi:10.1101/gad.1255805

Bantele, S.C.S., Lisby, M., Pfander, B., 2019. Quantitative sensing and signalling of single-stranded DNA during the DNA damage response. Nat Commun 10, 944. doi:10.1038/s41467-019-08889-5

Bartkova, J., Rezaei, N., Liontos, M., Karakaidos, P., Kletsas, D., Issaeva, N., Vassiliou, L.-V.F., Kolettas, E., Niforou, K., Zoumpourlis, V.C., Takaoka, M., Nakagawa, H., Tort, F., Fugger, K., Johansson, F., Sehested, M., Andersen, C.L., Dyrskjot, L., Ørntoft, T., Lukas, J., Kittas, C., Helleday, T., Halazonetis, T.D., Bartek, J., Gorgoulis, V.G., 2006. Oncogene-induced senescence is part of the tumorigenesis barrier imposed by DNA damage checkpoints. Nature 444, 633–637. doi:10.1038/nature05268

Bell, S.P., Labib, K., 2016. Chromosome Duplication in Saccharomyces cerevisiae. Genetics 203, 1027–1067. doi:10.1534/genetics.115.186452

Benaglia, T., Chauveau, D., Hunter, D.R., Young, D.S., 2009. mixtools: An R Package for Analyzing Finite Mixture Models. Journal of Statistical Software 32, 1–29. doi:10.18637/jss.v032.i06

Bittmann, J., Grigaitis, R., Galanti, L., Amarell, S., Wilfling, F., Matos, J., Pfander, B., 2020. An advanced cell cycle tag toolbox reveals principles underlying temporal control of structure-selective nucleases. Elife 9. doi:10.7554/eLife.52459

Blastyák, A., Pintér, L., Unk, I., Prakash, L., Prakash, S., Haracska, L., 2007. Yeast Rad5 protein required for postreplication repair has a DNA helicase activity specific for replication fork regression. Mol. Cell 28, 167–175. doi:10.1016/j.molcel.2007.07.030

Bleichert, F., 2019. Mechanisms of replication origin licensing: a structural perspective. Current Opinion in Structural Biology 59, 195–204. doi:10.1016/j.sbi.2019.08.007

Bui, D.T., Li, J.J., 2019. DNA Re-replication Is Susceptible to Nucleotide Level Mutagenesis. Genetics genetics.302194.2019–92. doi:10.1534/genetics.119.302194

Chabes, A., Domkin, V., Thelander, L., 1999. Yeast Sml1, a protein inhibitor of ribonucleotide reductase. Journal of Biological Chemistry 274, 36679–36683. doi:10.1074/jbc.274.51.36679

Chabes, A., Stillman, B., 2007. Constitutively high dNTP concentration inhibits cell cycle progression and the DNA damage checkpoint in yeast Saccharomyces cerevisiae. Proc Natl Acad Sci USA 104, 1183–1188. doi:10.1073/pnas.0610585104

Champeris Tsaniras, S., Kanellakis, N., Symeonidou, I.E., Nikolopoulou, P., Lygerou, Z., Taraviras, S., 2014. Licensing of DNA replication, cancer, pluripotency and differentiation: an interlinked world? Seminars in Cell and Developmental Biology 30, 174–180. doi:10.1016/j.semcdb.2014.03.013

Cheng, L., Collyer, T., Hardy, C.F., 1999. Cell cycle regulation of DNA replication initiator factor Dbf4p. Mol. Cell. Biol. 19, 4270–4278. doi:10.1128/mcb.19.6.4270

Cherry, J.M., Hong, E.L., Amundsen, C., Balakrishnan, R., Binkley, G., Chan, E.T., Christie, K.R., Costanzo, M.C., Dwight, S.S., Engel, S.R., Fisk, D.G., Hirschman, J.E., Hitz, B.C., Karra, K., Krieger, C.J., Miyasato, S.R., Nash, R.S., Park, J., Skrzypek, M.S., Simison, M., Weng, S., Wong, E.D., 2012. Saccharomyces Genome Database: the genomics resource of budding yeast. Nucleic Acids Research 40, D700–5. doi:10.1093/nar/gkr1029

Costantino, L., Sotiriou, S.K., Rantala, J.K., Magin, S., Mladenov, E., Helleday, T., Haber, J.E., Iliakis, G., Kallioniemi, O.P., Halazonetis, T.D., 2014. Break-Induced Replication Repair of Damaged Forks Induces Genomic Duplications in Human Cells. Science 343, 88–91. doi:10.1126/science.1243211

Cox, J., Mann, M., 2008. MaxQuant enables high peptide identification rates, individualized p.p.b.-range mass accuracies and proteome-wide protein quantification. Nat. Biotechnol. 26, 1367–1372. doi:10.1038/nbt.1511

Davé, A., Cooley, C., Garg, M., Bianchi, A., 2014. Protein phosphatase 1 recruitment by Rif1 regulates DNA replication origin firing by counteracting DDK activity. Cell Rep 7, 53–61. doi:10.1016/j.celrep.2014.02.019

Davidson, I.F., Li, A., Blow, J.J., 2006. Deregulated replication licensing causes DNA fragmentation consistent with head-to-tail fork collision. Mol. Cell 24, 433–443. doi:10.1016/j.molcel.2006.09.010

Davis, A.P., Symington, L.S., 2004. RAD51-dependent break-induced replication in yeast. Mol. Cell. Biol. 24, 2344–2351. doi:10.1128/MCB.24.6.2344-2351.2004

Drury, L.S., Diffley, J.F.X., 2009. Factors affecting the diversity of DNA replication licensing control in eukaryotes. Curr. Biol. 19, 530–535. doi:10.1016/j.cub.2009.02.034

Drury, L.S., Perkins, G., Diffley, J.F., 2000. The cyclin-dependent kinase Cdc28p regulates distinct modes of Cdc6p proteolysis during the budding yeast cell cycle. Current Biology 10, 231–240. doi:10.1016/s0960-9822(00)00355-9

Engel, S.R., Dietrich, F.S., Fisk, D.G., Binkley, G., Balakrishnan, R., Costanzo, M.C., Dwight, S.S., Hitz, B.C., Karra, K., Nash, R.S., Weng, S., Wong, E.D., Lloyd, P., Skrzypek, M.S., Miyasato, S.R., Simison, M., Cherry, J.M., 2014. The reference genome sequence of Saccharomyces cerevisiae: then and now. G3 (Bethesda) 4, 389–398. doi:10.1534/g3.113.008995

Ferreira, M.F., Santocanale, C., Drury, L.S., Diffley, J.F., 2000. Dbf4p, an essential S phase-promoting factor, is targeted for degradation by the anaphase-promoting complex. Mol. Cell. Biol. 20, 242–248. doi:10.1128/MCB.20.1.242-248.2000

Finn, K.J., Li, J.J., 2013. Single-Stranded Annealing Induced by Re-Initiation of Replication Origins Provides a Novel and Efficient Mechanism for Generating Copy Number Expansion via Non-Allelic Homologous Recombination. PLoS Genet. 9, e1003192–15. doi:10.1371/journal.pgen.1003192

Foiani, M., Lucchini, G., Plevani, P., 1997. The DNA polymerase alpha-primase complex couples DNA replication, cell-cycle progression and DNA-damage response. Trends Biochem. Sci. 22, 424–427. doi:10.1016/s0968-0004(97)01109-2

Forey, R., Poveda, A., Sharma, S., Barthe, A., Padioleau, I., Renard, C., Lambert, R., Skrzypczak, M., Ginalski, K., Lengronne, A., Chabes, A., Pardo, B., Pasero, P., 2020. Mec1 Is Activated at the Onset of Normal S Phase by Low-dNTP Pools Impeding DNA Replication. Mol. Cell 78, 1–20. doi:10.1016/j.molcel.2020.02.021

Fu, H., Redon, C.E., Thakur, B.L., Utani, K., Sebastian, R., Jang, S.-M., Gross, J.M., Mosavarpour, S., Marks, A.B., Zhuang, S.Z., Lazar, S.B., Rao, M., Mencer, S.T., Baris, A.M., Pongor, L.S., Aladjem, M.I., 2021. Dynamics of replication origin over-activation. Nat Commun 12, 3448. doi:10.1038/s41467-021-23835-0

Gambus, A., Jones, R.C., Sanchez-Diaz, A., Kanemaki, M., van Deursen, F., Edmondson, R.D., Labib, K., 2006. GINS maintains association of Cdc45 with MCM in replisome progression complexes at eukaryotic DNA replication forks. Nat. Cell Biol. 8, 358–366. doi:10.1038/ncb1382

Green, B.M., Finn, K.J., Li, J.J., 2010. Loss of DNA replication control is a potent inducer of gene amplification. Science 329, 943–946. doi:10.1126/science.1190966

Green, B.M., Li, J.J., 2005. Loss of rereplication control in Saccharomyces cerevisiae results in extensive DNA damage. Mol. Biol. Cell 16, 421–432. doi:10.1091/mbc.E04-09-0833

Green, B.M., Morreale, R.J., Ozaydin, B., Derisi, J.L., Li, J.J., 2006. Genome-wide mapping of DNA synthesis in Saccharomyces cerevisiae reveals that mechanisms preventing reinitiation of DNA replication are not redundant. Mol. Biol. Cell 17, 2401–2414. doi:10.1091/mbc.E05-11-1043

Guarino, E., Salguero, I., Kearsey, S.E., 2014. Cellular regulation of ribonucleotide reductase in eukaryotes. Seminars in Cell and Developmental Biology 30, 97–103. doi:10.1016/j.semcdb.2014.03.030

Hall, B.M., Ma, C.-X., Liang, P., Singh, K.K., 2009. Fluctuation analysis CalculatOR: a web tool for the determination of mutation rate using Luria-Delbruck fluctuation analysis. Bioinformatics 25, 1564–1565. doi:10.1093/bioinformatics/btp253

Hall, J.R., Lee, H.O., Bunker, B.D., Dorn, E.S., Rogers, G.C., Duronio, R.J., Cook, J.G., 2008. Cdt1 and Cdc6 are destabilized by rereplication-induced DNA damage. Journal of Biological Chemistry 283, 25356–25363. doi:10.1074/jbc.M802667200

Hanlon, S.L., Li, J.J., 2015. Re-replication of a Centromere Induces Chromosomal Instability and Aneuploidy. PLoS Genet. 11, e1005039–30. doi:10.1371/journal.pgen.1005039

Hennion, M., Arbona, J.-M., Lacroix, L., Cruaud, C., Theulot, B., Tallec, B.L., Proux, F., Wu, X., Novikova, E., Engelen, S., Lemainque, A., Audit, B., Hyrien, O., 2020. FORK-seq: replication landscape of the Saccharomyces cerevisiae genome by nanopore sequencing. Genome Biol. 21, 125. doi:10.1186/s13059-020-02013-3

Hiraga, S.-I., Alvino, G.M., Chang, F., Lian, H.-Y., Sridhar, A., Kubota, T., Brewer, B.J., Weinreich, M., Raghuraman, M.K., Donaldson, A.D., 2014. Rif1 controls DNA replication by directing Protein Phosphatase 1 to reverse Cdc7-mediated phosphorylation of the MCM complex. Genes Dev. 28, 372–383. doi:10.1101/gad.231258.113

Holt, L.J., Tuch, B.B., Villén, J., Johnson, A.D., Gygi, S.P., Morgan, D.O., 2009. Global analysis of Cdk1 substrate phosphorylation sites provides insights into evolution. Science 325, 1682–1686. doi:10.1126/science.1172867

Ira, G., Haber, J.E., 2002. Characterization of RAD51-independent break-induced replication that acts preferentially with short homologous sequences. Mol. Cell. Biol. 22, 6384–6392. doi:10.1128/MCB.22.18.6384-6392.2002

Jang, S.-M., Zhang, Y., Utani, K., Fu, H., Redon, C.E., Marks, A.B., Smith, O.K., Redmond, C.J., Baris, A.M., Tulchinsky, D.A., Aladjem, M.I., 2018. The replication initiation determinant protein (RepID) modulates replication by recruiting CUL4 to chromatin. Nat Commun 9, 2782–13. doi:10.1038/s41467-018-05177-6

Janke, C., Magiera, M.M., Rathfelder, N., Taxis, C., Reber, S., Maekawa, H., Moreno-Borchart, A., Doenges, G., Schwob, E., Schiebel, E., Knop, M., 2004. A versatile toolbox for PCR-based tagging of yeast genes: new fluorescent proteins, more markers and promoter substitution cassettes. Yeast 21, 947–962. doi:10.1002/yea.1142

Kim, J.C., Nordman, J., Xie, F., Kashevsky, H., Eng, T., Li, S., MacAlpine, D.M., Orr-Weaver, T.L., 2011. Integrative analysis of gene amplification in Drosophila follicle cells: parameters of origin activation and repression. Genes Dev. 25, 1384–1398. doi:10.1101/gad.2043111

Kim, U.J., Han, M., Kayne, P., Grunstein, M., 1988. Effects of histone H4 depletion on the cell cycle and transcription of Saccharomyces cerevisiae. EMBO J. 7, 2211–2219. doi:10.1002/j.1460-2075.1988.tb03060.x

Klotz-Noack, K., McIntosh, D., Schurch, N., Pratt, N., Blow, J.J., 2012. Re-replication induced by geminin depletion occurs from G2 and is enhanced by checkpoint activation. Journal of Cell Science 125, 2436–2445. doi:10.1242/jcs.100883

Kramara, J., Osia, B., Malkova, A., 2018. Break-Induced Replication: The Where, The Why, and The How. Trends in Genetics 34, 518–531. doi:10.1016/j.tig.2018.04.002

Kurat, C.F., Lambert, J.-P., Petschnigg, J., Friesen, H., Pawson, T., Rosebrock, A., Gingras, A.-C., Fillingham, J., Andrews, B., 2014. Cell cycle-regulated oscillator coordinates core histone gene transcription through histone acetylation. Proc Natl Acad Sci USA 111, 14124–14129. doi:10.1073/pnas.1414024111

Lee, Y.D., Wang, J., Stubbe, J., Elledge, S.J., 2008. Dif1 Is a DNA-Damage-Regulated Facilitator of Nuclear Import for Ribonucleotide Reductase. Mol. Cell 32, 70–80. doi:10.1016/j.molcel.2008.08.018

Li, H., Durbin, R., 2010. Fast and accurate long-read alignment with Burrows-Wheeler transform. Bioinformatics 26, 589–595. doi:10.1093/bioinformatics/btp698

Li, H., Handsaker, B., Wysoker, A., Fennell, T., Ruan, J., Homer, N., Marth, G., Abecasis, G., Durbin, R., 1000 Genome Project Data Processing Subgroup, 2009. The Sequence Alignment/Map format and SAMtools. Bioinformatics 25, 2078–2079. doi:10.1093/bioinformatics/btp352

Lopez-Delisle, L., Rabbani, L., Wolff, J., Bhardwaj, V., Backofen, R., Grüning, B., Ramírez, F., Manke, T., 2021. pyGenomeTracks: reproducible plots for multivariate genomic datasets. Bioinformatics 37, 422–423. doi:10.1093/bioinformatics/btaa692

Lopez-Mosqueda, J., Maas, N.L., Jonsson, Z.O., DeFazio-Eli, L.G., Wohlschlegel, J., Toczyski, D.P., 2010. Damage-induced phosphorylation of Sld3 is important to block late origin firing. Nature 467, 479–483. doi:10.1038/nature09377

Macheret, M., Halazonetis, T.D., 2018. Intragenic origins due to short G1 phases underlie oncogene-induced DNA replication stress. Nature 555, 112–116. doi:10.1038/nature25507

Machida, Y.J., Dutta, A., 2007. The APC/C inhibitor, Emi1, is essential for prevention of rereplication. Genes Dev. 21, 184–194. doi:10.1101/gad.1495007

Maiorano, D., Krasinska, L., Lutzmann, M., Méchali, M., 2005. Recombinant Cdt1 Induces Rereplication of G2 Nuclei in Xenopus Egg Extracts. Current Biology 15, 146–153. doi:10.1016/j.cub.2004.12.002

Mantiero, D., Mackenzie, A., Donaldson, A., Zegerman, P., 2011. Limiting replication initiation factors execute the temporal programme of origin firing in budding yeast. EMBO J. 30, 4805–4814. doi:10.1038/emboj.2011.404

Marzluff, W.F., Duronio, R.J., 2002. Histone mRNA expression: multiple levels of cell cycle regulation and important developmental consequences. Curr. Opin. Cell Biol. 14, 692–699. doi:10.1016/s0955-0674(02)00387-3

Mattarocci, S., Shyian, M., Lemmens, L., Damay, P., Altintas, D.M., Shi, T., Bartholomew, C.R., Thomä, N.H., Hardy, C.F.J., Shore, D., 2014. Rif1 Controls DNA Replication Timing in Yeast through the PP1 Phosphatase Glc7. Cell Rep 7, 62–69. doi:10.1016/j.celrep.2014.03.010

McGarry, T.J., Kirschner, M.W., 1998. Geminin, an inhibitor of DNA replication, is degraded during mitosis. Cell 93, 1043–1053. doi:10.1016/s0092-8674(00)81209-x

Melixetian, M., Ballabeni, A., Masiero, L., Gasparini, P., Zamponi, R., Bartek, J., Lukas, J., Helin, K., 2004. Loss of Geminin induces rereplication in the presence of functional p53. J. Cell Biol. 165, 473–482. doi:10.1083/jcb.200403106

Mendiratta, S., Gatto, A., Almouzni, G., 2019. Histone supply: Multitiered regulation ensures chromatin dynamics throughout the cell cycle. The Journal of Cell Biology 218, 39–54. doi:10.1083/jcb.201807179

Menzel, J., Tatman, P., Black, J.C., 2020. Isolation and analysis of rereplicated DNA by Rerep-Seq. Nucleic Acids Research 355, eaah6317–12. doi:10.1093/nar/gkaa197

Minca, E.C., Kowalski, D., 2010. Multiple Rad5 activities mediate sister chromatid recombination to bypass DNA damage at stalled replication forks. Mol. Cell 38, 649–661. doi:10.1016/j.molcel.2010.03.020

Moiseeva, T.N., Bakkenist, C.J., 2018. Regulation of the initiation of DNA replication in human cells. DNA Repair 72, 99–106. doi:10.1016/j.dnarep.2018.09.003

Morawska, M., Ulrich, H.D., 2013. An expanded tool kit for the auxin-inducible degron system in budding yeast. Yeast 30, 341–351. doi:10.1002/yea.2967

Mughal, M.J., Mahadevappa, R., Kwok, H.F., 2019. DNA replication licensing proteins: Saints and sinners in cancer. Semin Cancer Biol 58, 11–21. doi:10.1016/j.semcancer.2018.11.009

Müller, C.A., Boemo, M.A., Spingardi, P., Kessler, B.M., Kriaucionis, S., Simpson, J.T., Nieduszynski, C.A., 2019. Capturing the dynamics of genome replication on individual ultra-long nanopore sequence reads. Nat. Methods 16, 429–436. doi:10.1038/s41592-019-0394-y

Muñoz, S., Búa, S., Rodríguez-Acebes, S., Megías, D., Ortega, S., de Martino, A., Méndez, J., 2017. In Vivo DNA Re-replication Elicits Lethal Tissue Dysplasias. Cell Rep 19, 928–938. doi:10.1016/j.celrep.2017.04.032

Natsume, T., Müller, C.A., Katou, Y., Retkute, R., Gierliński, M., Araki, H., Blow, J.J., Shirahige, K., Nieduszynski, C.A., Tanaka, T.U., 2013. Kinetochores coordinate pericentromeric cohesion and early DNA replication by Cdc7-Dbf4 kinase recruitment. Mol. Cell 50, 661–674. doi:10.1016/j.molcel.2013.05.011

Neelsen, K.J., Zanini, I.M.Y., Mijic, S., Herrador, R., Zellweger, R., Ray Chaudhuri, A., Creavin, K.D., Blow, J.J., Lopes, M., 2013. Deregulated origin licensing leads to chromosomal breaks by rereplication of a gapped DNA template. Genes Dev. 27, 2537–2542. doi:10.1101/gad.226373.113

Nguyen, V.Q., Li, J.J., 2001. Cyclin-dependent kinases prevent DNA re-replication through multiple mechanisms. Nature 411, 1068–1073. doi:10.1038/35082600

Nordman, J.T., Kozhevnikova, E.N., Verrijzer, C.P., Pindyurin, A.V., Andreyeva, E.N., Shloma, V.V., Zhimulev, I.F., Orr-Weaver, T.L., 2014. DNA copy-number control through inhibition of replication fork progression. Cell Rep 9, 841–849. doi:10.1016/j.celrep.2014.10.005

Oshiro, G., Owens, J.C., Shellman, Y., Sclafani, R.A., Li, J.J., 1999. Cell cycle control of Cdc7p kinase activity through regulation of Dbf4p stability. Mol. Cell. Biol. 19, 4888–4896. doi:10.1128/MCB.19.7.4888

Peritore, M., Reußwig, K.-U., Bantele, S.C.S., Straub, T., Pfander, B., 2021. Strand-specific ChIP-seq at DNA breaks distinguishes ssDNA versus dsDNA binding and refutes single-stranded nucleosomes. Mol. Cell 1–27. doi:10.1016/j.molcel.2021.02.005

Petropoulos, M., Tsaniras, S.C., Taraviras, S., Lygerou, Z., 2019. Replication Licensing Aberrations, Replication Stress, and Genomic Instability. Trends Biochem. Sci. 1–13. doi:10.1016/j.tibs.2019.03.011

Pfander, B., Diffley, J.F.X., 2011. Dpb11 coordinates Mec1 kinase activation with cell cycle-regulated Rad9 recruitment. EMBO J. 30, 4897–4907. doi:10.1038/emboj.2011.345

Porcella, S.Y., Koussa, N.C., Tang, C.P., Kramer, D.N., Srivastava, P., Smith, D.J., 2020. Separable, Ctf4-mediated recruitment of DNA Polymerase α for initiation of DNA synthesis at replication origins and lagging-strand priming during replication elongation. PLoS Genet. 16, e1008755–23. doi:10.1371/journal.pgen.1008755

Putnam, C.D., Kolodner, R.D., 2010. Determination of gross chromosomal rearrangement rates. Cold Spring Harb Protoc 2010, pdb.prot5492. doi:10.1101/pdb.prot5492

Raghuraman, M.K., Winzeler, E.A., Collingwood, D., Hunt, S., Wodicka, L., Conway, A., Lockhart, D.J., Davis, R.W., Brewer, B.J., Fangman, W.L., 2001. Replication dynamics of the yeast genome. Science 294, 115–121. doi:10.1126/science.294.5540.115

Ramírez, F., Ryan, D.P., Grüning, B., Bhardwaj, V., Kilpert, F., Richter, A.S., Heyne, S., Dündar, F., Manke, T., 2016. deepTools2: a next generation web server for deep-sequencing data analysis. Nucleic Acids Research 44, W160–5. doi:10.1093/nar/gkw257

Reußwig, K.-U., Pfander, B., 2019. Control of Eukaryotic DNA Replication Initiation-Mechanisms to Ensure Smooth Transitions. Genes 10, 99. doi:10.3390/genes10020099

Reußwig, K.-U., Zimmermann, F., Galanti, L., Pfander, B., 2016. Robust Replication Control Is Generated by Temporal Gaps between Licensing and Firing Phases and Depends on Degradation of Firing Factor Sld2. Cell Rep 17, 556–569. doi:10.1016/j.celrep.2016.09.013

Richardson, C.D., Li, J.J., 2014. Regulatory Mechanisms That Prevent Re-initiation of DNA Replication Can Be Locally Modulated at Origins by Nearby Sequence Elements. PLoS Genet. 10, e1004358–19. doi:10.1371/journal.pgen.1004358

Rothstein, R.J., 1983. One-step gene disruption in yeast. Meth. Enzymol. 101, 202–211. doi:10.1016/0076-6879(83)01015-0

Santocanale, C., Diffley, J.F., 1998. A Mec1- and Rad53-dependent checkpoint controls late-firing origins of DNA replication. Nature 395, 615–618. doi:10.1038/27001

Schub, O., Rohaly, G., Smith, R.W., Schneider, A., Dehde, S., Dornreiter, I., Nasheuer, H.P., 2001. Multiple phosphorylation sites of DNA polymerase alpha-primase cooperate to regulate the initiation of DNA replication in vitro. Journal of Biological Chemistry 276, 38076–38083. doi:10.1074/jbc.M104975200

Shirahige, K., Hori, Y., Shiraishi, K., Yamashita, M., Takahashi, K., Obuse, C., Tsurimoto, T., Yoshikawa, H., 1998. Regulation of DNA-replication origins during cell-cycle progression. Nature 395, 618–621. doi:10.1038/27007

Siddiqui, K., On, K.F., Diffley, J.F.X., 2013. Regulating DNA replication in eukarya. Cold Spring Harb Perspect Biol 5. doi:10.1101/cshperspect.a012930

Siow, C.C., Nieduszynska, S.R., Muller, C.A., Nieduszynski, C.A., 2011. OriDB, the DNA replication origin database updated and extended. Nucleic Acids Research 40, D682–D686. doi:10.1093/nar/gkr1091

Tada, S., Li, A., Maiorano, D., Méchali, M., Blow, J.J., 2001. Repression of origin assembly in metaphase depends on inhibition of RLF-B/Cdt1 by geminin. Nat. Cell Biol. 3, 107–113. doi:10.1038/35055000

Tak, Y.-S., Tanaka, Y., Endo, S., Kamimura, Y., Araki, H., 2006. A CDK-catalysed regulatory phosphorylation for formation of the DNA replication complex Sld2-Dpb11. EMBO J. 25, 1987–1996. doi:10.1038/sj.emboj.7601075

Talarek, N., Petit, J., Gueydon, E., Schwob, E., 2015. EdU Incorporation for FACS and Microscopy Analysis of DNA Replication in Budding Yeast. Methods Mol. Biol. 1300, 105–112. doi:10.1007/978-1-4939-2596-4_7

Tanaka, S., 2021. Construction of Tight Conditional Mutants Using the Improved Auxin-Inducible Degron (iAID) Method in the Budding Yeast Saccharomyces cerevisiae. Methods Mol. Biol. 2196, 15–26. doi:10.1007/978-1-0716-0868-5_2

Tanaka, S., Araki, H., 2013. Helicase activation and establishment of replication forks at chromosomal origins of replication. Cold Spring Harb Perspect Biol 5, a010371. doi:10.1101/cshperspect.a010371

Tanaka, S., Araki, H., 2011. Multiple regulatory mechanisms to inhibit untimely initiation of DNA replication are important for stable genome maintenance. PLoS Genet. 7, e1002136. doi:10.1371/journal.pgen.1002136

Tanaka, S., Miyazawa-Onami, M., Iida, T., Araki, H., 2015. iAID: an improved auxin-inducible degron system for the construction of a “tight” conditional mutant in the budding yeast Saccharomyces cerevisiae. Yeast 32, 567–581. doi:10.1002/yea.3080

Tanaka, S., Nakato, R., Katou, Y., Shirahige, K., Araki, H., 2011. Origin Association of Sld3, Sld7,and Cdc45 Proteins Is a Key Stepfor Determination of Origin-Firing Timing. Current Biology 21, 2055–2063. doi:10.1016/j.cub.2011.11.038

Tanaka, S., Umemori, T., Hirai, K., Muramatsu, S., Kamimura, Y., Araki, H., 2007. CDK-dependent phosphorylation of Sld2 and Sld3 initiates DNA replication in budding yeast. Nature 445, 328–332. doi:10.1038/nature05465

Tanny, R.E., MacAlpine, D.M., Blitzblau, H.G., Bell, S.P., 2006. Genome-wide analysis of re-replication reveals inhibitory controls that target multiple stages of replication initiation. Mol. Biol. Cell 17, 2415–2423. doi:10.1091/mbc.E05-11-1037

Vassilev, A., DePamphilis, M.L., 2017. Links between DNA Replication, Stem Cells and Cancer. Genes 8, 45. doi:10.3390/genes8020045

Vrtis, K.B., Dewar, J.M., Chistol, G., Wu, R.A., Graham, T.G.W., Walter, J.C., 2021. Single-strand DNA breaks cause replisome disassembly. Mol. Cell 81, 1309–1318.e6. doi:10.1016/j.molcel.2020.12.039

Walter, D., Hoffmann, S., Komseli, E.-S., Rappsilber, J., Gorgoulis, V., Sørensen, C.S., 2016. SCF(Cyclin F)-dependent degradation of CDC6 suppresses DNA re-replication. Nat Commun 7, 10530–10. doi:10.1038/ncomms10530

Wang, W., Klein, K.N., Proesmans, K., Yang, H., Marchal, C., Zhu, X., Borrman, T., Hastie, A., Weng, Z., Bechhoefer, J., Chen, C.-L., Gilbert, D.M., Rhind, N., 2021. Genome-wide mapping of human DNA replication by optical replication mapping supports a stochastic model of eukaryotic replication. Mol. Cell 81, 2975–2988.e6. doi:10.1016/j.molcel.2021.05.024

Wohlschlegel, J.A., Dwyer, B.T., Dhar, S.K., Cvetic, C., Walter, J.C., Dutta, A., 2000. Inhibition of eukaryotic DNA replication by geminin binding to Cdt1. Science 290, 2309–2312. doi:10.1126/science.290.5500.2309

Zegerman, P., Diffley, J.F.X., 2010. Checkpoint-dependent inhibition of DNA replication initiation by Sld3 and Dbf4 phosphorylation. Nature 467, 474–478. doi:10.1038/nature09373

Zegerman, P., Diffley, J.F.X., 2007. Phosphorylation of Sld2 and Sld3 by cyclin-dependent kinases promotes DNA replication in budding yeast. Nature 445, 281–285. doi:10.1038/nature05432

Zhao, X., Muller, E.G., Rothstein, R., 1998. A suppressor of two essential checkpoint genes identifies a novel protein that negatively affects dNTP pools. Mol. Cell 2, 329–340. doi:10.1016/s1097-2765(00)80277-4

Zhou, Y., Pozo, P.N., Oh, S., Stone, H.M., Cook, J.G., 2020. Distinct and sequential re-replication barriers ensure precise genome duplication. PLoS Genet. 16, e1008988–24. doi:10.1371/journal.pgen.1008988

Zhu, W., Chen, Y., Dutta, A., 2004. Rereplication by Depletion of Geminin Is Seen Regardless of p53 Status and Activates a G2/M Checkpoint. Mol. Cell. Biol. 24, 7140–7150. doi:10.1128/MCB.24.16.7140-7150.2004

